# The human Y and inactive X chromosomes similarly modulate autosomal gene expression

**DOI:** 10.1101/2023.06.05.543763

**Authors:** Adrianna K. San Roman, Helen Skaletsky, Alexander K. Godfrey, Neha V. Bokil, Levi Teitz, Isani Singh, Laura V. Blanton, Daniel W. Bellott, Tatyana Pyntikova, Julian Lange, Natalia Koutseva, Jennifer F. Hughes, Laura Brown, Sidaly Phou, Ashley Buscetta, Paul Kruszka, Nicole Banks, Amalia Dutra, Evgenia Pak, Patricia C. Lasutschinkow, Colleen Keen, Shanlee M. Davis, Angela E. Lin, Nicole R. Tartaglia, Carole Samango-Sprouse, Maximilian Muenke, David C. Page

**Affiliations:** Whitehead Institute; Cambridge, MA 02142, USA; Department of Biology, Massachusetts Institute of Technology; Cambridge, MA 02139, USA; Howard Hughes Medical Institute, Whitehead Institute; Cambridge, MA 02142, USA; Harvard Medical School, Boston, MA 02115, USA; Medical Genetics Branch, National Human Genome Research Institute, National Institutes of Health, Bethesda; MD 20892, USA; Eunice Kennedy Shriver National Institute of Child Health and Human Development, National Institutes of Health; Bethesda, MD 20892 USA; Cytogenetics and Microscopy Core, National Human Genome Research Institute, National Institutes of Health; Bethesda, MD 20892 USA; Focus Foundation, Davidsonville, MD 21035, USA; Department of Pediatrics, University of Colorado School of Medicine, Aurora, CO 80045, USA; Medical Genetics, Massachusetts General for Children, Boston, MA 02114, USA; Department of Pediatrics, Harvard Medical School, Boston, MA 02115, USA; Developmental Pediatrics, eXtraOrdinarY Kids Program, Children’s Hospital Colorado, Aurora, CO 80011, USA; Department of Pediatrics, George Washington University, Washington, DC 20052, USA; Department of Human and Molecular Genetics, Florida International University, Miami, FL 33199, USA

**Keywords:** Sex chromosomes, sex differences, X chromosome inactivation, aneuploidy, Turner syndrome, Klinefelter syndrome, gene expression, transcription factors, CRISPR

## Abstract

Somatic cells of human males and females have 45 chromosomes in common, including the “active” X chromosome. In males the 46^th^ chromosome is a Y; in females it is an “inactive” X (Xi). Through linear modeling of autosomal gene expression in cells from individuals with zero to three Xi and zero to four Y chromosomes, we found that Xi and Y impact autosomal expression broadly and with remarkably similar effects. Studying sex-chromosome structural anomalies, promoters of Xi- and Y-responsive genes, and CRISPR inhibition, we traced part of this shared effect to homologous transcription factors – *ZFX* and *ZFY* – encoded by Chr X and Y. This demonstrates sex-shared mechanisms by which Xi and Y modulate autosomal expression. Combined with earlier analyses of sex-linked gene expression, our studies show that 21% of all genes expressed in lymphoblastoid cells or fibroblasts change expression significantly in response to Xi or Y chromosomes.

## INTRODUCTION

The human X and Y chromosomes evolved from ordinary autosomes over the course of the last 200 million years.^1,2^ Today, these homologous chromosomes – typically two X chromosomes in females, and one X and one Y chromosome in males – comprise the oldest, most massive variation in the human genome.

Until recently, this chromosome-scale variation was understood to be of little direct consequence outside the reproductive tract, where the decisive roles of sex chromosome constitution are well established. The reasoning was straightforward: the second X chromosome in 46,XX cells is condensed and transcriptionally attenuated through the process of X chromosome inactivation (XCI),^3,4^ and the Y chromosome carries relatively few genes, many of which are not expressed outside the testes.^5,6^

Despite this logic, observations in recent decades have suggested more expansive roles for the second (“inactive”) X chromosome in human female cells,^7,8^ and even for the Y chromosome in male somatic cells.^9–11^ In theory, differing sex chromosome complements in somatic cells could help explain the abundant male-female differences in autosomal gene expression that are seen throughout the human body.^12–14^ Still, it is not known whether these differences in autosomal transcriptomes are driven by sex chromosome constitution, sex steroid hormones, or other biological, behavioral, or environmental factors.

Diploid human somatic cells invariably have one active X chromosome (Xa); most also have an inactive X (Xi) or a Y chromosome, and some have additional Xi or Y chromosomes. We previously modeled expression of X- or Y-linked genes as a function of Xi or Y copy number in cells cultured from individuals with various sex chromosome constitutions: one to four copies of chromosome (Chr) X (zero to three copies of Xi) and zero to four copies of Chr Y.^15^ Expression of 38% of X-linked genes changed with additional copies of Xi, and expression of nearly all Y-linked genes increased with additional copies of Chr Y. These Xi- and Y-driven changes were remarkably modular, with successive Xi or Y chromosomes having quantitatively similar effects, facilitating linear modeling. By incorporating allele-specific analyses, we determined that the Chr X gene expression changes are due to a combination of 1) expression from Xi alleles and 2) modulation of Xa transcript levels by Xi, in trans.

Given that Xi modulates Xa genes in trans, the question arises whether Xi also modulates expression of autosomal genes, potentially via similar mechanisms. A related question is whether Chr Y modulates autosomal gene expression. Based on our recent success in linear modeling of sex-chromosome gene expression,^15^ we now adapt and expand that modeling to measure how variation in the number of X or Y chromosomes affects autosomal transcription in cultured human cells. We quantified the impact of variation in both Xi and Y chromosome copy number on autosomal transcripts, explored the mechanism(s) responsible for those effects, and discovered that a conserved pair of transcription factors expressed from Xi and Y contribute to modulation of gene expression on autosomes and Xa. An unanticipated finding is that the autosomal impacts of Xi and Y chromosomes are remarkably similar.

## RESULTS

### Thousands of autosomal genes respond to Chr X and Chr Y copy number

We examined the effects of Chr X and Chr Y copy number on autosomal gene expression using our recently reported RNA-seq dataset from 106 lymphoblastoid cell lines (LCLs) and 99 primary dermal fibroblast cultures.^15^ These lines and cultures derive from 176 individuals with one to four X chromosomes and zero to four Y chromosomes (**Fig. 1A, Table S1**). For each expressed autosomal gene, including protein-coding and long non-coding RNA genes with at least one transcript per million in either 46,XX or 46,XY samples, we fit a linear regression model in DESeq2^16^ to estimate the log_2_ fold change in expression per copy of Chr X or Y (**Methods**).

**Figure 1.**
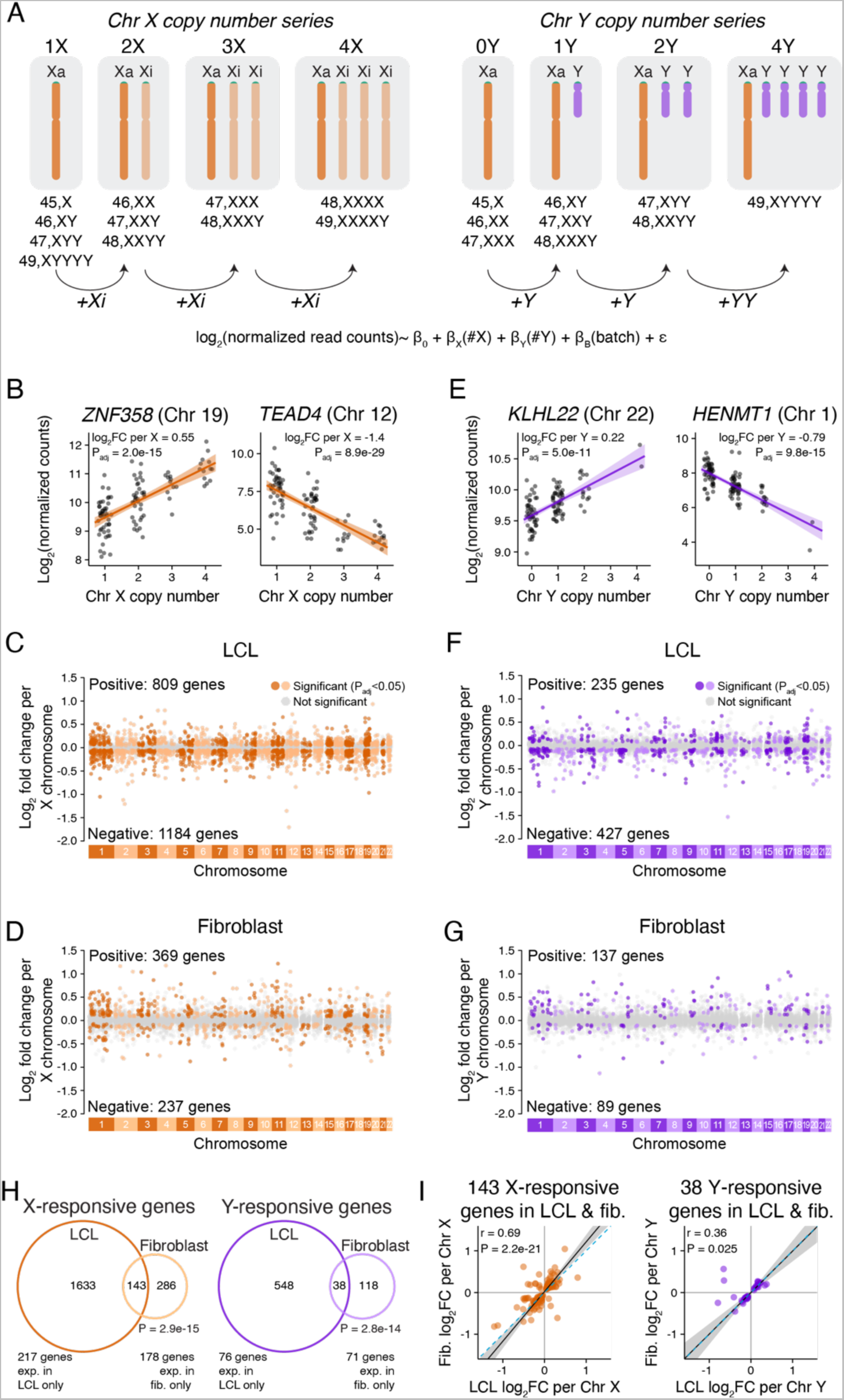
Genome-wide response to Chr X or Y copy number is distinct in two cell types. **(A)** RNA-seq data from LCLs and fibroblasts spanning a range of Chr X and Y copy numbers were analyzed using linear regression, modeling log_2_ expression as a function of Chr X copy number, Chr Y copy number, and batch. **(B,E)** Examples of individual autosomal genes that significantly respond to Chr X (B) or Chr Y (E) copy number in LCLs. Each point represents the expression level in an individual sample with the indicated number of X or Y chromosomes. The regression lines and confidence intervals, log_2_ fold change per copy of Chr X or Y, and adjusted P values (P_adj_) from linear regressions are indicated. **(C,D,F,G)** Log_2_ fold change per copy of Chr X or Y for 11,034 expressed autosomal genes in LCLs and 12,002 expressed autosomal genes in fibroblasts, plotted by chromosomal location. Genes in grey do not significantly change in response to X or Y copy number, while genes in dark or light colors (odd- or even-numbered chromosomes, respectively) represent significantly Chr X- (C, D) or Chr Y-responsive (F, G) genes. **(H)** Venn diagrams showing the overlap of genes that significantly respond to X or Y copy number (P_adj_<0.05) in LCLs and fibroblasts. Genes expressed in both cell types were included in the Venn diagrams, and genes with cell-type specific expression are noted below. P values, hypergeometric test. **(I)** Scatterplots of genes that are Chr X- or Y-responsive in both LCLs and fibroblasts show mostly similar log_2_ fold change (log_2_FC) values across cell types. Black line and grey shading, weighted Deming regression and 95% confidence interval; blue dashed line, identity (X=Y) line. Pearson correlation coefficients and P values are indicated.

This analysis revealed that Chr X and Chr Y copy number had widespread effects on autosomal expression. The magnitudes of individual gene expression changes per Chr X or Y were modest – with most changes less than 1.5-fold – but many genes were affected. There were 1,993 significantly (adjusted P value [P_adj_]<0.05) X-responsive genes in LCLs – comprising 18% of expressed autosomal genes – and 606 in fibroblasts – 5% of expressed autosomal genes (**Fig. 1B-D;** full results in **Table S2**). Chr Y copy number affected somewhat fewer autosomal genes: 662 Chr Y-responsive genes in LCLs – 6% of expressed autosomal genes – and 226 in fibroblasts – 1.9% of expressed autosomal genes (**Fig. 1E-G, Table S2**). Gene ontology analyses of the Chr X- or Y-responsive genes yielded few significantly enriched functional categories (**Methods, Fig. S1**). We conclude that variation in Chr X and Y copy numbers significantly and broadly alters autosomal gene expression.

We considered and tested various interpretations of these findings. First, we asked whether X- or Y-responsive autosomal genes were statistical artifacts of enhanced sex-chromosomal expression altering the genome-wide distribution of sampled read counts. To this end, we renormalized the data and refit the linear models using only autosomal genes, obtaining virtually identical results (**Fig. S2**).

Second, we tested whether there was any impact of modeling gene expression as a function of both variables – Chr X copy number and Chr Y copy number – together compared to models in which we used subsets of samples to vary either Chr X or Chr Y copy number individually. We found that the full model was highly correlated genome-wide with models of the individual contributions of Chr X and Y copy number (**Fig. S3**). Moreover, by conducting the Chr X analysis in samples with and without Y chromosomes, we found that the autosomal effects of Xi copy number are comparable in males and females (**Fig. S3**).

Third, we tested the impact of sample size on the number of significantly Chr X- or Y-responsive genes detected in our analysis to ask whether we have saturated the analysis with our current sample set. Using a bootstrapping analysis, we found that sequencing cell lines from additional individuals would lead to discovery of more sex-chromosome-responsive genes, but they would likely have small effect sizes (**Fig. S4**).

Finally, and because individuals across our dataset have differing hormonal profiles that might be correlated with Chr X or Y copy number, we evaluated whether a memory of differential hormonal exposures in the body prior to tissue sampling might explain the results. Most X- or Y-responsive autosomal genes did not overlap with direct target genes for estrogen or androgen receptors (**Methods, Table S3, Fig. S5**).^17,18^ There was a modest enrichment of androgen receptor targets among Y-responsive genes in fibroblasts, but their effects were not correlated (Spearman r=0.034, P= 0.9; **Fig. S5D**). We conclude that the impact of Chr X or Y copy number is independent of sex hormones. In sum, thousands of autosomal genes respond significantly to Chr X or Y copy number, with more genes responding to X than to Y, and more in LCLs than in fibroblasts.

### Response to Chr X or Chr Y copy number is mostly cell-type-specific

We investigated whether individual autosomal genes responded to Chr X or Chr Y copy number in LCLs only, in fibroblasts only, or in both cell types. Across autosomal genes expressed in both cell types, we observed a weak correlation between the response to Chr X and Chr Y (**Fig. S6**). Most significantly responsive genes were responsive in only one of the two cell types, but among genes expressed in both cell types, the overlap was greater than expected by chance: 34% of X-responsive and 25% of Y-responsive genes in fibroblasts were also X- or Y-responsive in LCLs (**Fig. 1H**). Most of these overlapping genes responded with the same directionality – rising (or falling) in response to Chr X or Y copy number in both LCLs and fibroblasts (**Fig. 1I**). Thus, some of the genome-wide response to sex-chromosome copy number is shared between the two cell types, but most is cell-type-specific.

### Chr Xi and Y elicit similar responses across autosomes and Xa

Next, we compared, within each cell type, the gene-by-gene autosomal responses to Chr X and to Chr Y. We observed highly correlated responses to Chr X and Y (**Fig. 2A**), such that additional copies of Xi and Y had similar effects across the genome. Restricting this analysis to autosomal genes with statistically significant responses to Chr X or Y copy number showed that >60% of Y-responsive genes were also X-responsive – 442 genes in LCLs and 136 in fibroblasts (**Fig. 2B**). These co-regulated genes’ responses to X and Y were highly concordant in polarity and magnitude, suggesting that the mechanisms governing these responses might be shared (**Fig. 2C, S7A**).

**Figure 2.**
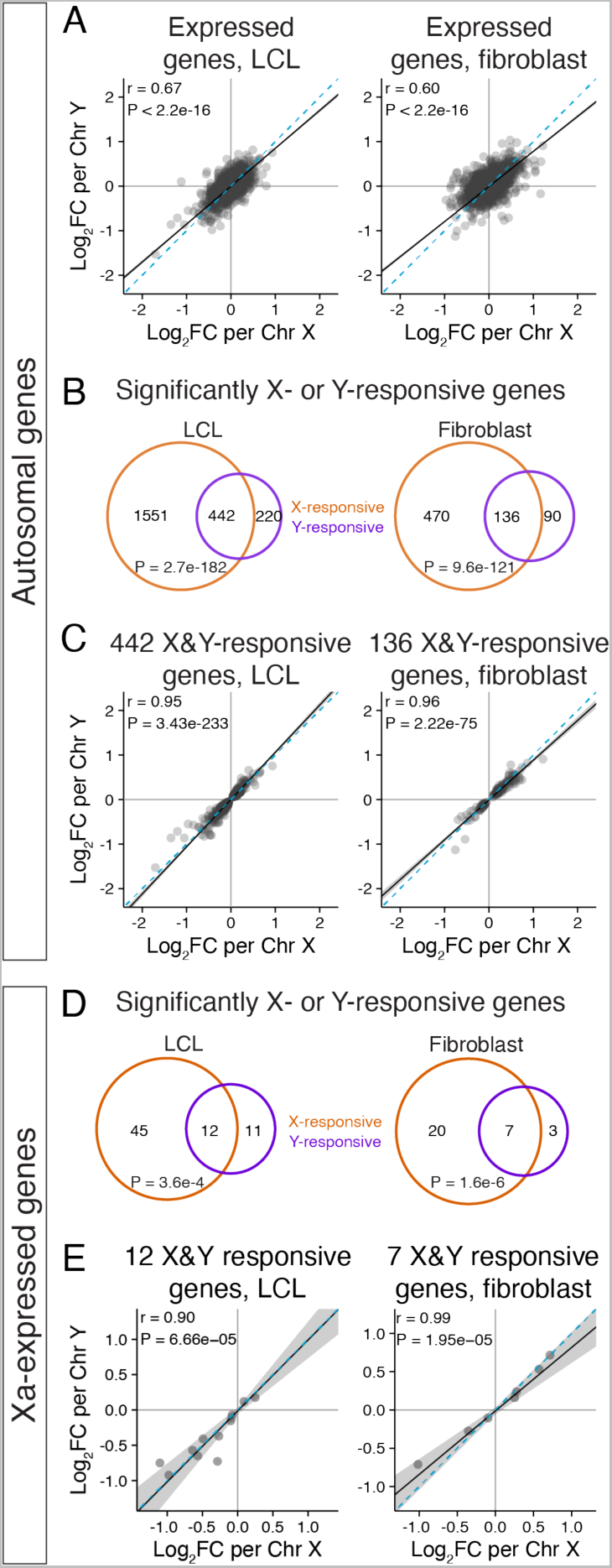
Chr X and Y copy number have similar genome-wide effects. **(A)** Scatterplots of all expressed autosomal genes in LCLs and fibroblasts show globally correlated response to Chr X and Chr Y copy number. **(B,D)** Venn diagrams of significantly (P_adj_<0.05) X- and Y-responsive autosomal (B) or Xa-expressed (D) genes reveal significant overlap. P values, hypergeometric test. **(C,E)** Scatterplots of the log_2_ fold change (log_2_FC) per Chr X vs Chr Y for significantly Chr X- and Y-responsive autosomal (C) or Xa-expressed (E) genes show correlated response. Black line and grey shading, weighted Deming regression and 95% confidence interval; blue dashed line, identity (X=Y) line; Pearson correlation coefficients and P values are indicated.

In a previous study, we showed that Xi copy number modulates gene expression from Xa in trans.^15^ In light of our current findings, we wondered whether Chr X and Chr Y copy number might have shared effects on Xa gene expression. We first conducted a new meta-analysis, compiling data from studies quantifying Xi expression to identify strictly “Xa-expressed” genes that are consistently “silenced” on Xi across studies (**Methods; Table S4**).^7,8,15, 19–22^ Using these annotations, we found that slightly larger proportions of Xa-expressed genes were impacted by Chr X or Y copy number than on the autosomes: 57 (20%) and 27 (8.6%) Xa-expressed genes were significantly responsive to Chr X copy number in LCLs and fibroblasts, respectively, and 23 (8.2%) and 10 (3.2%) were responsive to Chr Y copy number (**Table S4**). Among these, 12 genes in LCLs and 7 genes in fibroblasts significantly responded to both Chr X and Y copy numbers (**Fig. 2D**). Like the autosomal genes, the responses of Xa-expressed genes to Chr X and Chr Y copy number were highly concordant in magnitude and polarity (**Fig. 2E, S7B**). This suggests that Xi and Y modulate Xa gene expression in trans through shared mechanisms.

To identify the mechanisms driving the genome-wide shared response to Chr X and Chr Y copy numbers, we considered and tested three hypotheses. We asked whether the observed expression changes reflect 1) a generic response to aneuploidy that is not specific to the sex chromosomes, 2) a “heterochromatin sink” created by additional sex chromosomes that pulls heterochromatin factors away from and thereby de-represses (activates) autosomal genes, or 3) the action of regulatory genes shared between the X and Y chromosomes. We will consider each of these hypotheses in turn.

### The shared response to Chr X and Chr Y copy number is not a generic response to aneuploidy

To test the first hypothesis, we analyzed RNA-seq data from LCLs with three copies of Chr 21 (47,XX+21 and 47,XY+21) and compared these with our data from 46,XX and 46,XY LCLs (**Fig. 3A, Table S1**).^15^ We employed linear modeling to estimate the log_2_ fold change in expression per additional Chr 21, controlling for sex chromosome constitution (46,XX or 46,XY) (**Methods**). Chr 21 had large effects across the genome: in addition to expression increases for Chr 21 genes (previously reported in San Roman *et al.*^15^), we identified 980 genes on other chromosomes (including X and Y) that were significantly Chr 21-responsive (**Fig. 3B-D, Table S5**).

**Figure 3.**
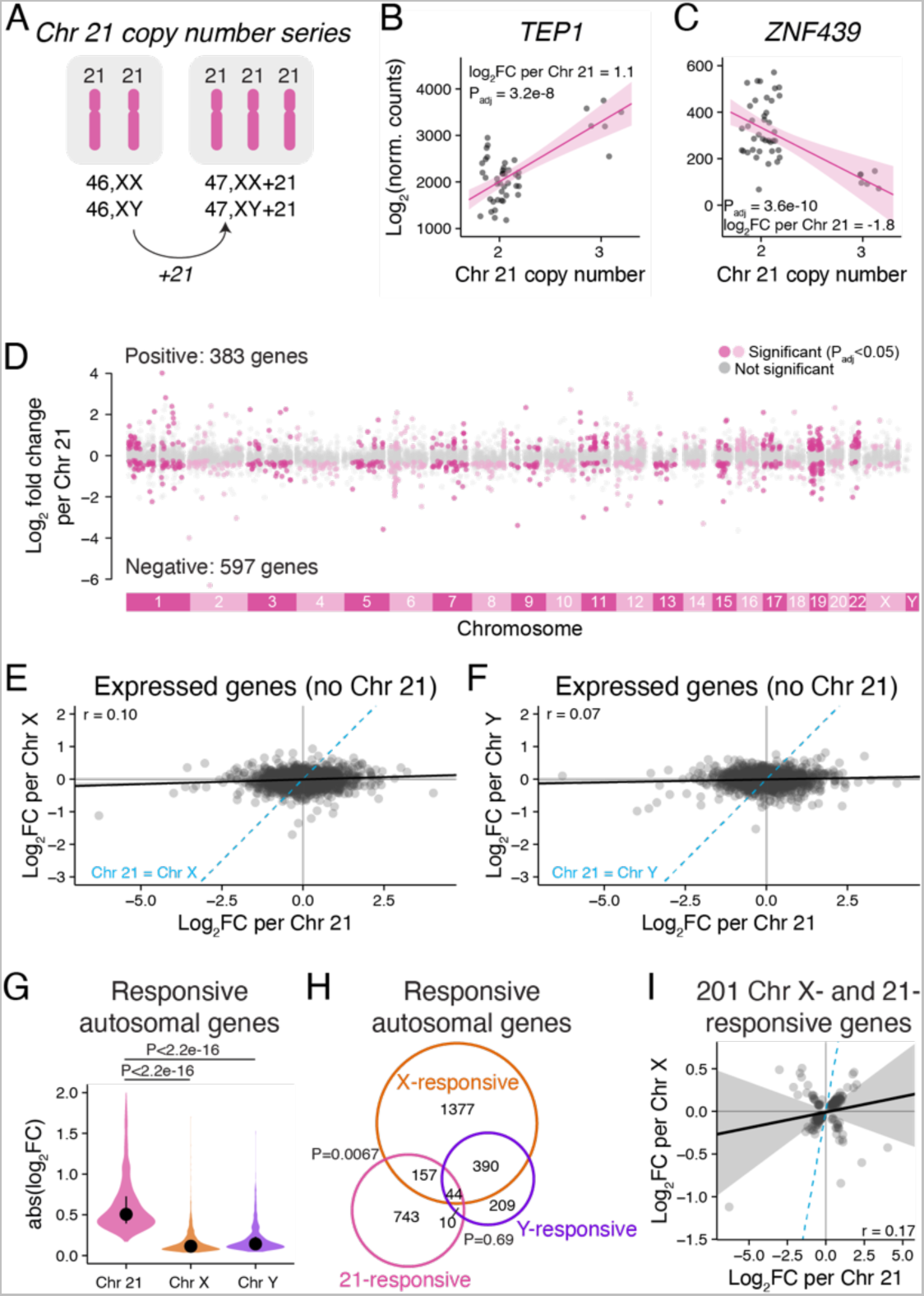
The shared response to Chr X and Y copy number is not a general response to aneuploidy. **(A)** RNA-seq data from LCLs with two or three copies of Chr 21 were analyzed using linear regression. **(B-C)** Scatterplots and regression lines with confidence intervals of individual genes showing expression changes in samples with two or three copies of Chr 21. The log_2_ fold change per Chr 21 and P_adj_ from linear regressions are indicated. **(D)** The log_2_ fold changes per Chr 21 across all chromosomes except for Chr 21 are plotted by chromosomal location. Genes in grey are expressed, but do not significantly change in response to Chr 21 copy number, while genes in dark or light pink (colored by every other chromosome for clarity) represent significantly Chr 21-responsive genes. **(E-F)** Scatterplot of all expressed autosomal genes, except Chr 21, comparing Chr 21 response to Chr X (E) or Chr Y (F) response. Blue dashed line, Chr 21 = Chr X (E) or Chr Y (F). **(G)** Violin plot of the absolute values of the log_2_ fold changes for significantly Chr 21, X, or Y-responsive genes. P values, Wilcoxon rank sum test. **(H)** Venn diagram of Chr X-, Chr Y- and Chr 21-responsive genes shows a significant overlap between genes that respond to Chr X and Chr 21 (P values, hypergeometric test). **(I)** Scatterplot of genes regulated by both Chr X and Chr 21 shows little correlation between their responses to X and to 21. All scatterplots: black line and grey shading, Deming regression and 95% confidence interval; blue dashed line, identity (X=Y) line; Pearson correlation coefficients are indicated.

We then compared the autosomal responses to Chr 21 copy number with responses to Chr X or Chr Y copy number. Across the autosomes, Chr 21-responsive genes generally displayed larger fold changes than did Chr X- or Y-responsive genes (**Fig. 3E-G**). Importantly, most X- or Y-responsive autosomal genes did not overlap with the Chr 21-responsive gene set (**Fig. 3H**). There was a modest overlap between Chr 21-responsive and Chr X-responsive genes, but the effects of Chr 21 and Chr X were weakly correlated (**Fig. 3I**). Moreover, none of the Xa-expressed genes that responded to both Chr X and Y copy number in LCLs were among the Chr 21-responsive Chr X genes (**Fig. S8**). We conclude that features unique to the sex chromosomes, rather than a generic aneuploidy response, explain autosomal (and Xa) gene expression changes in response to Chr X and Y copy number.

### The response to Chr Y copy number is not due to heterochromatin sinks

“Heterochromatin sink” effects are well documented in *Drosophila*, where Y chromosomes with varying amounts of heterochromatin affect expression of autosomal genes adjacent to heterochromatic regions – a phenomenon known as position-effect variegation.^23^ We studied two kinds of human variation to test whether Chr Y heterochromatin impacts autosomal gene expression: 1) common polymorphisms in heterochromatin length on the long arm of Chr Y (Yq),^24^ and 2) rare structurally variant Y chromosomes.

To quantify Yq heterochromatin, we mapped whole-genome sequencing data from 1225 male samples from the 1000 Genomes Project^25^ to *DYZ1*, a repetitive sequence specific to the Yq heterochromatic region.^6^ The average depth of coverage of the *DYZ1* sequence ranged from 23.8 to 896.5 reads (after normalization to depth of coverage of a single-copy region of the Y chromosome), with a median of 202.5 (**Fig. S9A; Table S6**). This variation in *DYZI* read depth was corroborated by fluorescent in situ hybridization (FISH) to metaphase chromosomes (**Fig. S9B; Table S6**). For 194 samples for which RNA-seq data was also available,^26^ we modeled gene expression as a function of *DYZ1* read depth, finding only three significantly differentially expressed genes (**Fig. 4A-B**). We conclude that variation in Yq heterochromatin length does not affect genome-wide expression.

**Figure 4.**
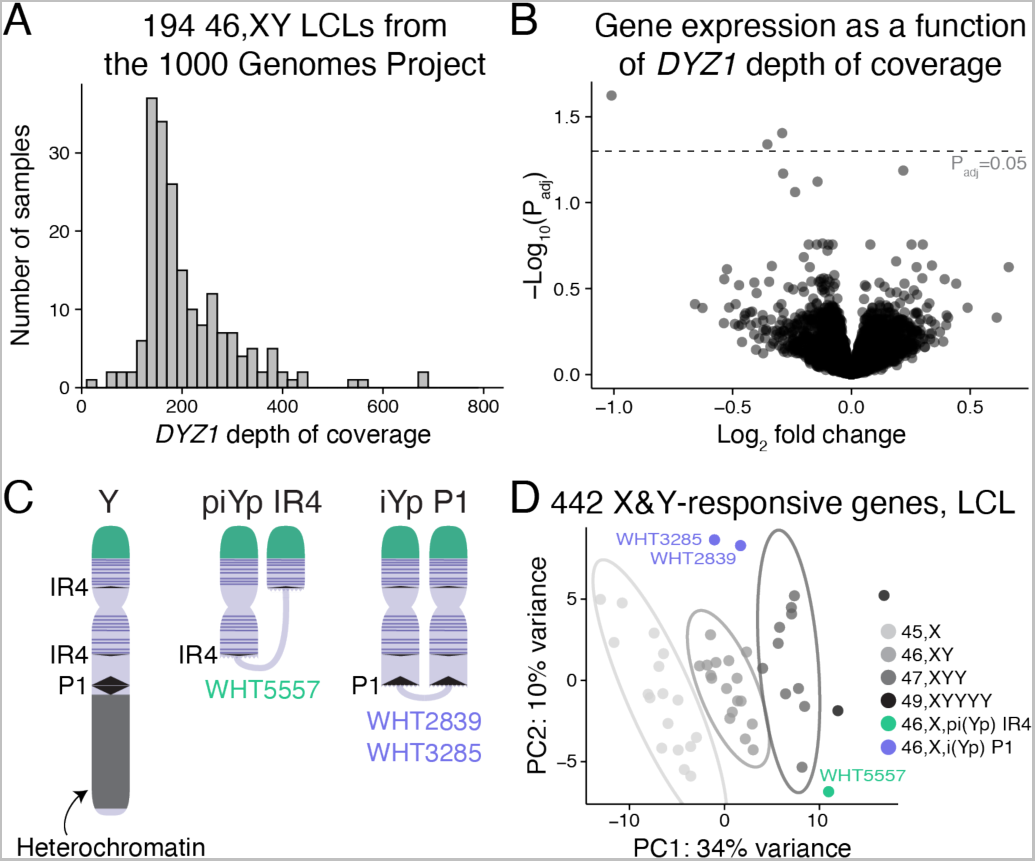
Heterochromatin sink model does not explain genes upregulated in response to Chr Y copy number. **(A)** Histogram of *DYZ1* depth of coverage for the 194 46,XY samples from the 1000 Genomes Project that have paired RNA-sequencing data. **(B)** Each point represents the magnitude and significance of expression change in response to *DYZ1* depth of coverage for expressed autosomal genes in LCLs. Only three genes are below the significance cutoff of P_adj_=0.05. **(C)** Schematic of the normal Y chromosome and two types of variant Y chromosomes that have recombined in repeated DNA regions to generate chromosomes missing the large heterochromatic region on the long arm and have two copies of many or all Chr Y genes expressed in somatic cells (purple horizontal lines). One Y pseudoisochromosome (piYp) resulted from recombination in an inverted repeat (IR4) on Yp and Yq, and a different Y isochromosome (iYp) resulted from recombination in a palindrome (P1) on Yq. **(D)** PCA plot showing separation of samples based on expression of the 442 autosomal genes that are X- and Y-responsive in LCLs. Ellipses represent 95% confidence intervals around the centroid of each karyotype group with at least three samples. The 46,X,i(Yp) and 46,X,pi(Yp) samples cluster away from 45,X samples and near those with two Y chromosomes (47,XYY).

We separated the effects of Chr Y genes from effects of Yq heterochromatin using LCLs from individuals with structurally variant Y “isochromosomes” in which Yq heterochromatin is deleted and a portion of the chromosome, including widely-expressed Chr Y genes on the short arm (Yp), is duplicated in “mirror image” orientation, resulting in higher Chr Y gene expression (**Fig. 4C, S10; Table S1**). If Yq heterochromatin contributes significantly to expression of shared X- and Y-responsive genes, we would expect cells missing this region to show expression like that of 45,X cells, which also lack Yq heterochromatin. If Chr Y genes are responsible, we would expect these samples to show expression like that of 47,XYY samples. Using principal component analysis of the 442 autosomal genes responsive to both Chr X and Y copy number in LCLs, we found that samples with variant Y chromosomes did not cluster with 45,X samples but were instead more similar to 47,XYY samples (**Fig. 4D**). Together, these experiments indicate that a heterochromatin sink does not explain the response of autosomal genes to Chr Y copy number, and therefore cannot explain the shared response to Chr X and Chr Y copy number.

### NPX-NPY gene pairs drive shared autosomal response to X and Y copy number

Two groups of genes shared between Chr X and Y could drive the shared autosomal response (**Fig. 5A**). First are genes in the shared “pseudoautosomal region” (PAR) on the distal short arms of Chr X and Y; expression of these genes increases linearly with sex chromosome copy number.^15^ Second are homologous gene pairs in the non-pseudoautosomal regions of Chr X (NPX) and Y (NPY), which increase in expression with X or Y copy number, respectively. These NPX-NPY gene pairs are remnants of the ~200-million-year evolutionary process that differentiated a pair of ordinary autosomes into Chr X, which retains 98% of the ancestral gene content, and Chr Y, which retains only 3%. The NPX-NPY pairs are involved in important cellular processes, including transcription, epigenetic regulation, and translation,^10^ and have diverged in sequence to varying degrees,^6^ possibly maintaining identical functions. Of the NPX-NPY pair genes, several may affect genome-wide expression, including the histone lysine demethylases *KDM6A/UTY* and *KDM5C/KDM5D*, RNA helicases *DDX3X/DDX3Y*, and transcription factors *ZFX/ZFY*. To decouple the effects of PAR versus NPX-NPY pair genes, we used RNA-seq to analyze cell lines from individuals with structurally variant X or Y chromosomes in which segments are deleted or duplicated.

**Figure 5.**
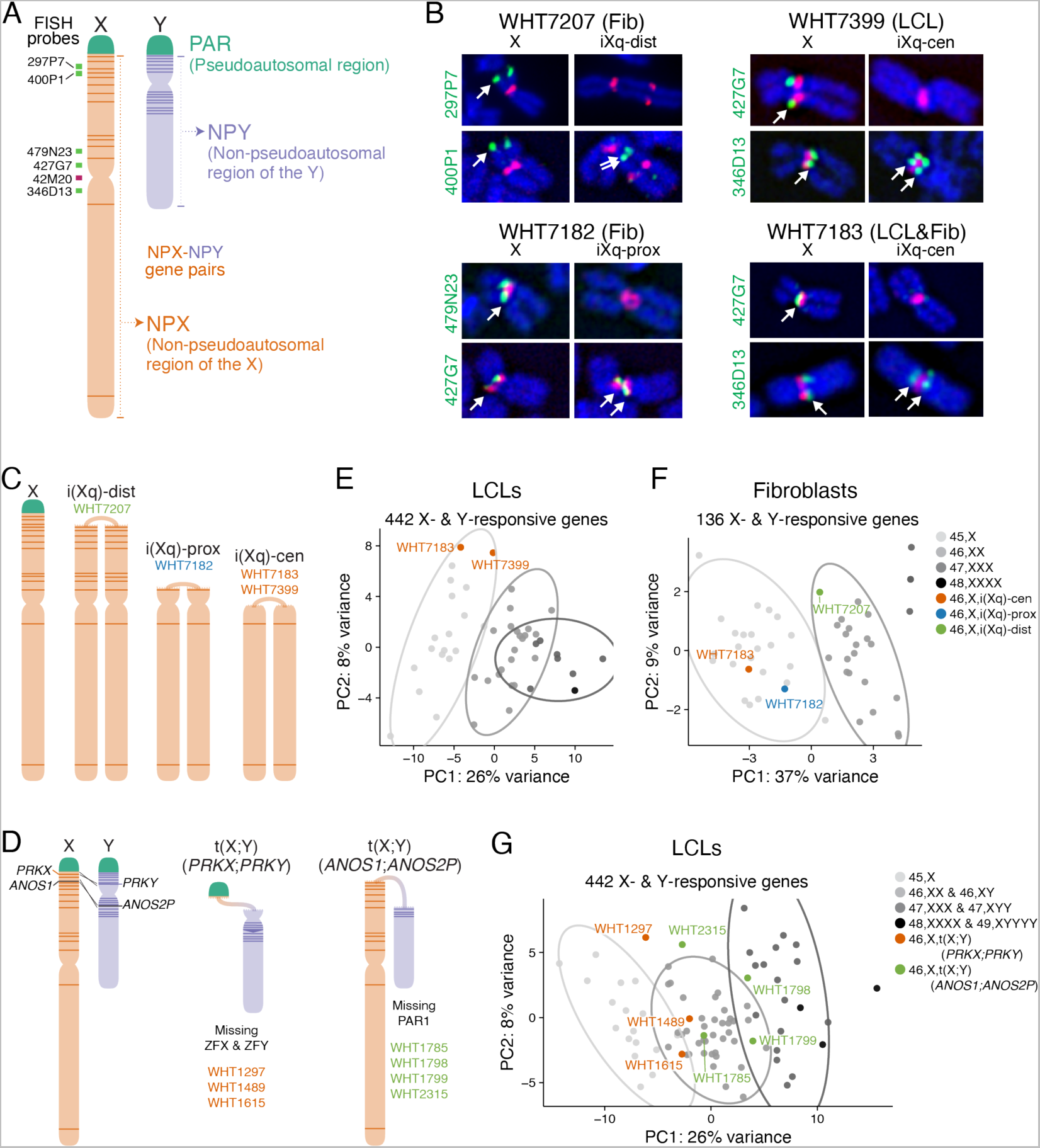
NPX-NPY pair genes on Xp (the short arm), but not PAR1 genes, drive the shared genome-wide response to X and Y chromosome copy number. **(A)** Anatomy of the sex chromosomes. Locations of FISH probes used in (B) are indicated. **(B)** DNA FISH on four 46,X,i(Xq) samples refined the sites of recombination. Fluorescent images of the normal X and iXq from the same cell are shown side-by-side, with probes across the chromosome (green) and the X centromere (red). Full FISH results are found in **Figure S11**. **(C)** Schematic of a normal Chr X and three types of X isochromosomes. Locations of NPX-NPY pair genes indicated by dark orange lines. **(D)** Schematic showing recombination between X and Y Chrs at *PRKX* and *PRKY*, or at *ANOS1* and *ANOS2P*, resulting in X-Y translocation products. **(E-F)** Principal component analysis (PCA) of autosomal X- and Y-responsive genes in LCL (E) or fibroblast (F) samples with one to three copies of Chr X or 46,X,i(Xq) structural variants. **(G)** PCA of autosomal X- and Y-responsive genes in LCL samples with one to five total sex chromosomes or with X-Y translocated chromosomes. Ellipses represent 95% confidence intervals around the centroid of each karyotype group with at least three samples.

We first investigated LCLs and fibroblasts from four individuals with structurally variant X chromosomes – isochromosomes -- containing two copies of the long arm (Xq) and missing all or part of the short arm (Xp) (**Table S1**). To map the recombination breakpoints for these X isochromosomes, we performed fluorescent in situ hybridization (FISH) using DNA probes for sequences across Chr X (**Fig. S11, Table S7**). We identified three different configurations (**Fig. 5B**): One individual’s isochromosome (referred to as iXq-dist) had a breakpoint in the distal short arm and is missing PAR1. Another individual’s isochromosome had a breakpoint in the proximal short arm (referred to as iXq-prox) and was further missing the NPX-NPY pair genes *ZFX, DDX3X,* and *KDM6A*. Finally, two individuals had breakpoints at the centromere (referred to as iXq-cen) and are further missing the NPX-NPY pair gene *KDM5C* (**Fig. 5C**; see **Table S7** for copy numbers of PAR1 and NPX-NPY pair genes in cell lines with structural variants). RNA-seq read counts were consistent with the breakpoints assigned by DNA FISH (**Fig. S12-14**).

The second group of cells was from individuals with one normal Chr X and an X-Y translocation product arising from recombination between highly similar genes in NPX and NPY, either *PRKX* and *PRKY* or *ANOS1 (KAL1)* and *ANOS2P (KALP)* (**Fig. 5D**).^27^ To identify samples with these X-Y translocations, we used a PCR-based screening strategy for Y chromosomes with breakpoints in regions encompassing *PRKY* and *ANOS2P* on our human cell line collection accumulated over three decades from individuals with karyotypic identification of an aberrant Y chromosome, discordance between sex chromosome constitution and sex phenotype, or spermatogenic failure (**Table S7**). We performed RNA-seq on three cell lines containing *PRKX-PRKY* translocated chromosomes, which retain PAR1 but have a deletion on distal Yp resulting in loss of the NPY genes *ZFY, SRY,* and *RPS4Y1* (**Table S7**). In addition, we performed RNA-seq on four cell lines with *ANOS1-ANOS2P* translocated chromosomes, which lack PAR1 but retain most NPX-NPY gene pairs (**Table S7**). RNA-seq results were again consistent with the chromosomal breakpoints mapped by PCR screening (**Fig. S12-14**).

To determine if the shared autosomal response to X and Y copy number is due to PAR1 or NPX-NPY genes, we compared gene expression in the cell lines with structurally variant sex chromosomes to those with sex chromosome aneuploidy, using principal component analysis of the shared Chr X- and Y-responsive autosomal genes in LCLs (442 genes) or fibroblasts (136 genes). If PAR1 genes drive the X-Y-shared autosomal response, we would expect all 46,X,i(Xq) and 46,X,t(X;Y)(*ANOS1-ANOS2P*) samples – which retain only one copy of PAR1 – to cluster with 45,X samples, and 46,X,t(X;Y)(*PRKX-PRKY*) samples *–* which retain two copies of PAR1 – to cluster with 46,XX and 46,XY samples. Instead, we found that the clustering of 46,X,i(Xq) samples depended on the copy number of NPX genes on Xp. For example, the 46,X,i(Xq-cen) and 46,X,i(Xq-prox) samples – with one copy of most or all NPX-NPY pair genes on Xp – clustered with the 45,X samples, while the 46,X,i(Xq-dist) sample – with three copies of most NPX-NPY pairs – clustered with the 46,XX samples (one might expect this sample to cluster with 47,XXX samples, but, due to mosaicism for a 45,X cell line (**Methods**), expression averages to 46,XX levels; **Fig. 5E-F**). Similarly, 46,X,t(X;Y)(*ANOS1-ANOS2P*) samples – with two copies of most NPX-NPY pairs – clustered with 46,XX samples, while 46,X,t(X;Y)(*PRKX-PRKY*) samples – with reduced copy number of only two NPX-NPY pairs: *ZFX/ZFY* and *RPS4X/RPS4Y* – clustered with 45,X samples (**Fig. 5G**). In all, these results indicate that NPX-NPY pairs, not PAR1 genes, drive the shared autosomal response to X and Y copy number in LCLs and fibroblasts. Among NPX-NPY pairs, only one pair both maps to Xp and is present in reduced copy number in 46,X,t(X;Y)(*PRKX-PRKY*) samples: *ZFX* and *ZFY*.

### *ZFX* and *ZFY* activate a shared transcriptional program genome-wide

To determine whether *ZFX* and *ZFY* mediate the genome-wide response to X or Y copy number, we searched for DNA motifs in promoters of X- or Y-responsive genes (**Methods**). Motifs matching previously-characterized ZFX DNA-binding signatures were enriched in genes that were positively X- or Y-responsive but not among genes that were negatively X- or Y-responsive; the highest enrichment was among genes positively responsive to *both* X and Y in LCLs (**Table S8**). Additionally, re-analysis of ENCODE data revealed that ZFX protein binding is enriched at promoters of genes positively responsive to X and Y in LCLs and fibroblasts (**Fig. S15, Table S9, Methods**). This aligns well with studies showing that ZFX is a transcriptional activator,^28–31^ and extends previous reports of ZFX motif enrichment in genes positively regulated by Chr X copy number identified using microarray data.^32,33^ The motif database we queried does not include a ZFY motif, but *ZFX* and *ZFY* are 93% identical in both DNA and predicted amino acid sequence^6^ and evidence indicates that ZFY occupies the same genomic sites as ZFX: their zinc-finger domains and genome-wide binding profiles are nearly identical.^34–36^

To directly assess the functions of ZFX and ZFY genome-wide, we knocked down *ZFX* in 46,XX fibroblasts (from three individuals) and *ZFX* and/or *ZFY* in 46,XY fibroblasts (from three individuals) using CRISPR interference (CRISPRi^37^; **Fig. 6A, Methods**), followed by RNA sequencing. To identify differentially expressed genes, we modeled gene expression as a function of the CRISPRi target gene (*ZFX* and/or *ZFY* versus control; **Methods**). These CRISPRi knockdowns were effective: in single knockdowns we observed an ~80% reduction of *ZFX* and >90% reduction of *ZFY* levels (**Fig. 6B**). The knockdowns were also specific: *ZFX* expression was not affected by *ZFY* knockdown, and vice-versa. In a separate experiment, double knockdowns were performed aiming for more modest but significant reductions of both *ZFX* (~40%) and *ZFY* (~55%).

**Figure 6.**
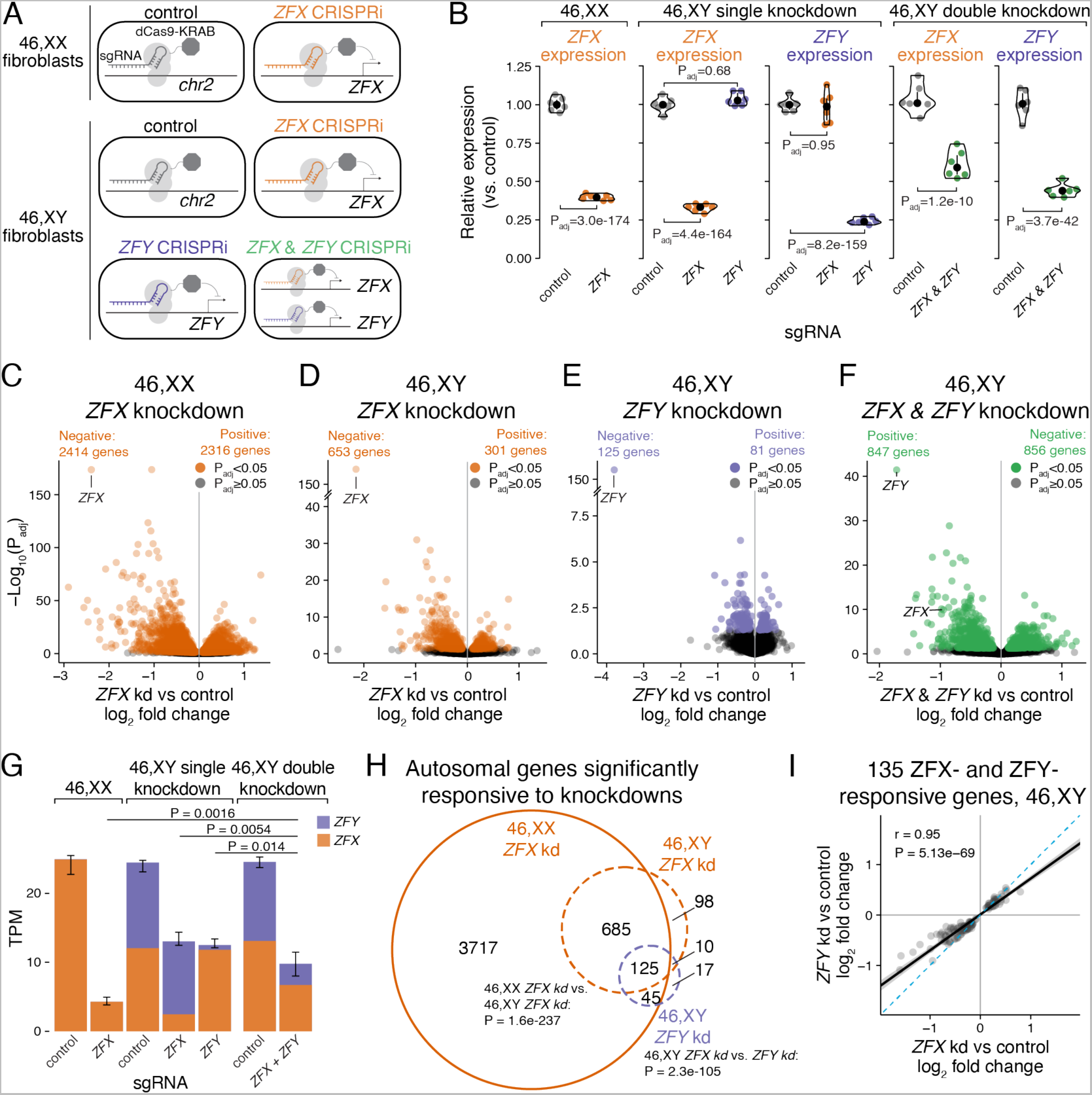
*ZFX* and *ZFY* activate a common set of genes. **(A)** Schematic of CRISPRi knockdown experiments. 46,XX and 46,XY fibroblasts from three individuals were transduced with dCas9-KRAB and sgRNAs directed against a control intergenic region on Chr 2, the promoter of *ZFX,* or the promoter of *ZFY* to block transcription. **(B)** Violin plots showing relative expression of *ZFX* or *ZFY* in cells transduced with *ZFX* and/or *ZFY* sgRNAs, compared to control sgRNAs. Log_2_FC and P_adj_ values from DESeq2 results. **(C-F)** Volcano plots showing effect size and significance of all expressed genes in *ZFX* and/or *ZFY* knockdowns versus controls. Numbers of significantly differentially expressed genes are indicated. **(G)** Stacked barplots showing median and interquartile ranges (whiskers) of cumulative expression in transcripts per million (TPM) of *ZFX* and *ZFY* in CRISPRi experiments. P values, t-tests. **(H)** Venn diagram of significantly differentially expressed autosomal genes upon knockdown of *ZFX* in XX or XY cells or upon knockdown of *ZFY* in XY cells. **(I)** Scatterplot of significantly *ZFX-* and *ZFY-*responsive autosomal genes in 46,XY cells comparing effects of *ZFX* versus *ZFY* knockdowns. Black line and grey shading, weighted Deming regression and 95% confidence interval; blue dashed line, identity (X=Y) line; Pearson correlation and P value are indicated.

The magnitude of the genome-wide effects varied among the knockdowns: in 46,XX cells, 4,638 genes were differentially expressed upon *ZFX* knockdown, while in 46,XY cells, 943 genes were differentially expressed upon *ZFX* knockdown, 206 genes upon *ZFY* knockdown, and 1,698 genes upon *ZFX* / *ZFY* double knockdown (**Fig. 6C-F, Table S10**). The knockdowns confirm that ZFX is a transcriptional activator: genes that decrease in expression with *ZFX* and/or *ZFY* knockdown in 46,XX and 46,XY cells were enriched for ZFX DNA binding motifs and protein binding at their promoters, compared to genes that were not significantly affected; conversely, genes that increased in expression were depleted for ZFX binding (**Fig. S16, Table S11**).

Noting that the genome-wide response to *ZFX* knockdown was markedly different in 46,XX and 46,XY cells, we hypothesized that expression of *ZFY* could play a role. Indeed, the magnitude of the genome-wide effects was inversely related to the cumulative level of *ZFX* and *ZFY* expression remaining after knockdown (**Fig. 6G**). Double-knockdown cells had lower total *ZFX+ZFY* levels compared to total *ZFX+ZFY* levels in either single knockdown, and they displayed a larger genome-wide effect, even though *ZFX* levels in the double knockdown were higher than in the single knockdowns. This suggests that *ZFX* and *ZFY* act in a mutually and cumulatively dose-dependent fashion, which we tested by comparing the responses to their single knockdowns. Among autosomal genes that responded to *ZFY* knockdown in 46,XY cells, 135 (69%) also responded to *ZFX* knockdown in 46,XY cells, and these responses were nearly perfectly correlated, although the genes displayed larger responses to *ZFX* than to *ZFY* knockdown (P=7.2×10^−8^, paired t-test; **Fig. 6H-I**). Additionally, all four Xa-expressed genes that significantly responded to *ZFY* knockdown also responded to *ZFX* knockdown in 46,XX and 46,XY cells (**Fig. S17**). We conclude that *ZFX* and *ZFY* activate the same genes, with *ZFX* being more potent than *ZFY*, and that the total levels of *ZFX*+*ZFY* impact thousands of genes across the genome.

### ZFX and ZFY are required for shared response to Chr X and Chr Y copy number

Finally, we asked whether *ZFX* or *ZFY* activity is required for the genome-wide response to Chr X and Chr Y copy number. Of the 136 autosomal genes responsive to both Chr X and Y copy number in fibroblasts, 109 (81%) responded to *ZFX* and/or *ZFY* knockdown, a significant enrichment (**Fig. 7A**). Among genes that responded significantly only to Chr X or only to Chr Y, 70% and 57%, respectively, were *ZFX* or *ZFY* responsive (**Fig. S18A**). On a gene-by-gene basis, the effects of *ZFX* or *ZFY* knockdown were negatively correlated with the response to Chr X or Chr Y copy number, consistent with ZFX and ZFY’s functions as transcriptional activators (**Fig. 7B, Fig. S18B)**. In all, the transcriptional changes upon *ZFX* or *ZFY* knockdown explained 19-40% of the effects for autosomal genes responsive to *either* Chr X or Y copy number, and 46-57% of the effects for autosomal genes responsive to *both* Chr X and Y copy number (**Fig. 7B, S18**).

**Figure 7.**
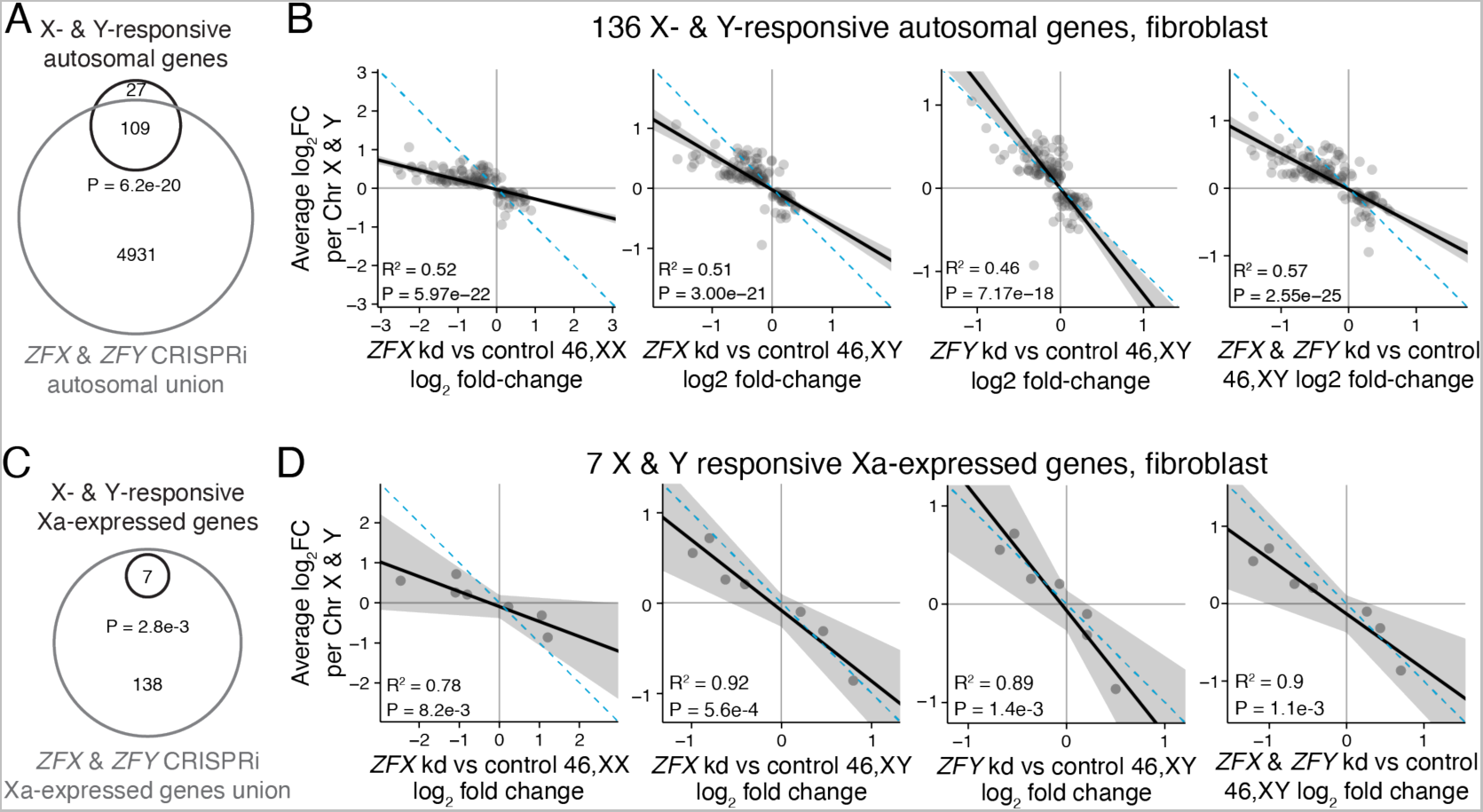
*ZFX* and *ZFY* are required for shared Chr X and Chr Y responsive transcriptional program. **(A, C)** Venn diagram of X- and Y-responsive autosomal (A) or Xa-expressed (C) genes in fibroblasts with union of all *ZFX*- and *ZFY*-responsive genes across the four CRISPRi experiments. P value, hypergeometric test. **(B, D)** Scatterplots comparing response to Chr X copy number and response to *ZFX* or *ZFY* knockdown for 136 autosomal genes (B) or 7 Xa-expressed genes (D) that are significantly X- and Y-responsive in fibroblasts. Black line and grey shading, weighted Deming regression and 95% confidence interval; blue dashed line, identity (X=Y) line; coefficients of determination and P values are indicated.

We obtained similar results for genes that are only expressed from Xa. Of the seven Xa-expressed genes significantly responsive to Chr X or Y number in fibroblasts, all were significantly responsive to *ZFX* and/or *ZFY* knockdown (**Fig. 7C**). For these seven Xa-expressed genes, the effects of *ZFX* and/or *ZFY* knockdown explain 78-92% of the effects of Chr X and Y copy number (**Fig. 7D**). In sum, the actions of *ZFX* and *ZFY* explain most of the shared response to Chr X and Chr Y copy number on Xa as well as on autosomes.

## DISCUSSION

Traditionally, the “gene-poor” Y and “inactive” X (Xi) chromosomes have not been considered prominent players in regulating the transcriptome of human somatic cells. Studies of these “sex chromosomes” have focused instead on their capacity to differentiate the reproductive cells and organs of males and females. Our study produced two surprising findings that challenge common understandings of these chromosomes. First, the Xi and Y chromosomes modulate expression of thousands of autosomal genes in somatic cells, as we demonstrated in transformed (lymphoblastoid) cells and primary cultures (skin fibroblasts). Second, the genome-wide impact of these two chromosomes is strikingly similar: many autosomal genes respond to Xi and Y by changing expression in the same direction and with correlated effect sizes. These shared effects do not reflect a generic aneuploidy response or a response to a heterochromatin sink, and they are not driven by pseudoautosomal genes. Instead, a pair of NPX- and NPY-encoded transcriptional activators, ZFX and ZFY, account for about half of the shared effects.

### 21% of autosomal genes expressed in LCLs or fibroblasts respond to Chr X or Y copy number

We began this study by quantitatively modeling autosomal gene expression in cells of two types cultured from 176 individuals with a wide range of sex chromosome constitutions: one to four X chromosomes (zero to three Xi chromosomes), and zero to four Y chromosomes (e.g., 45,X to 49,XXXXY and 49,XYYYY). Incorporating these diverse sex chromosome constitutions into a single linear model (**Fig. 1**) yielded advantages over earlier studies that compared transcriptomes in pairwise fashion across a circumscribed set of sex chromosome constitutions: most frequently comparing 46,XX with 46,XY; 45,X with 46,XX; and 46,XY with 47,XXY.^12–14,32,33,38–43^ This linear model provided the power required to detect and quantify increases or decreases in expression of individual autosomal genes as a function of Chr X copy number, and as a function of Chr Y copy number. Eighteen percent of LCL-expressed autosomal genes were significantly responsive to Chr X copy number, and 6% of such genes were responsive to Chr Y copy number (**Fig. 1**). In fibroblasts, 5% of expressed autosomal genes were significantly X-responsive, and 2% were Y-responsive. Merging these observations, we find that, of 13,126 autosomal genes expressed in LCLs and/or fibroblasts, 2,722 (20.7%) are significantly responsive to Chr X and/or Chr Y copy number in one or both cell types (**Table S12**). Additionally, of 335 Xa-expressed genes in LCLs and/or fibroblasts, 88 (26.3%) are significantly responsive to Chr X and/or Chr Y copy number in one or both cell types (**Table S12**). As a group, these Xi- or Y-responsive genes -- 2,722 autosomal genes and 88 Xa-expressed genes -- are widely dispersed across functional categories, with only a weak enrichment for functional categories related to metabolism (**Fig. S19**).

Importantly, we observed these effects of Xi and Y copy number even when minimizing the differentiating influence of sex steroid hormones by culturing cells under identical conditions. Moreover, we found that the effects of Xi copy number on autosomal gene expression are comparable in cells derived from phenotypic males and females. We conclude that Xi effects on autosomal gene expression are independent of the presence of the Y chromosome or the gonadal sex of the individual donor.

### Shared effects on the transcriptome reflect the common autosomal ancestry of X and Y chromosomes

The Y and X chromosomes play starkly different roles in the differentiation and function of the human reproductive tract.^44^ In cultured somatic cells, however, we observed that the Y and Xi chromosomes have broadly similar and highly correlated effects on the transcriptome of all other chromosomes, including the Xa chromosome found in somatic cells of all individuals, regardless of their biological sex.

We had not anticipated these similar effects of Xi and Y on the global transcriptome, but they can be explained in light of the evolution of the human X and Y chromosomes from a pair of autosomes over the past 200 million years.^1,2^ The human X and Y chromosomes originated not as “sex chromosomes” but as ordinary autosomes, identical in sequence and, in all likelihood, largely identical in their functions in male and female somatic cells. The genes of these ancestral autosomes included global mediators and regulators of gene expression unrelated to sex differentiation – mediators and regulators that survived the differentiation of the X and Y chromosomes and remain expressed today on both Y and Xi.^10,45^

Indeed, as shown here, one of these global regulators – surviving today as *ZFX* on the human X chromosome and its homolog *ZFY* on the human Y chromosome – accounts for about half of the shared effect of Y and Xi on global gene expression. Combining the results reported here with prior discoveries, we propose the following mechanistic model: The single Xa chromosome found in all somatic cells (regardless of sex) contributes a relatively invariant level of ZFX protein; each Xi chromosome in the cell adds a smaller amount of ZFX protein, and each Y chromosome adds a similar quantity of ZFY protein. In trans, ZFX and ZFY proteins act additively, and functionally interchangeably, in modulating transcript levels for thousands of autosomal genes (and Xa genes).

The details and generalizability of this model should be tested in diverse somatic cell types. However, our findings are sufficient to state that the shared autosomal ancestry of the Xi and Y chromosomes continues to shape the functional roles of both chromosomes. Recognizing that *ZFX* (and *ZFY*) account for about half of the Xi and Y chromosomes’ effects on the autosomal transcriptome in LCLs and skin fibroblasts, we encourage investigators to explore the possibility that other ancestral NPX-NPY gene pairs contribute mechanistically to sex-shared functions of the Xi and Y chromosomes.

### The challenge: distinguishing between the shared and differentiating roles of Xi and Y in human somatic cells

Our findings demonstrate that human Xi and Y chromosomes perform shared roles in modulating the autosomal (and Xa) transcriptome in somatic cells. However, myriad differences in autosomal gene expression in human somatic tissues (outside the reproductive tract) are found when comparing 46,XY males to 46,XX females.^12–14,38,39^ More research is required to determine the degree to which these transcriptomic differences represent sex differences in cellular composition, sex steroid hormones, or cell-autonomous roles of the Xi and Y chromosomes. The task of searching for and molecularly substantiating sex-differentiating, cell-autonomous roles of individual Xi- or Y-expressed genes in somatic tissues, at the molecular level, will be complicated by the breadth and scale of the sex-shared, cell-autonomous roles described here.

### Limitations of this study

We found a genome-wide response to Chr X and Y copy number in both LCLs and fibroblasts, but most of the autosomal genes that responded were X- and/or Y-responsive in only one of the two cell types. Thus, these autosomal “responders” will likely differ based on the epigenetic context of the cell or tissue type of interest, and this should be taken into consideration when comparing the gene lists here to metrics derived in other tissues. We note that this same cell-type variation could occur for androgen or estrogen receptor direct target genes, and so the absence of an overlap with X- and Y-responsive genes could reflect different targets in cancer cell lines versus LCLs and fibroblasts. The heterochromatin sink hypothesis was tested (and excluded) by studying naturally occurring variation in the Y chromosome; this hypothesis was not directly excluded for the X chromosome, where we could not genetically or experimentally separate the possible effects of Xi heterochromatin from the effects of Xi gene expression.

## Acknowledgements

We thank members of the Page laboratory for discussions, Sahin Naqvi for advice on RNA-sequencing and analysis, Jorge Adarme and Susan Tocio for laboratory support, and the Whitehead Institute Genome Technology Core facility for library preparation and sequencing. We thank the individuals who contributed tissue samples for their participation.

## Funding

National Institutes of Health grant F32HD091966 (AKSR)

National Institutes of Health grant U01HG0007587 (DCP and MM)

National Institutes of Health grant K23HD092588 (SMD)

National Science Foundation Graduate Research Fellowship grant 1745302 (NVB)

Simons Foundation Autism Research Initiative – Collaboration Award (DCP, JFH, DWB)

Lallage Feazel Wall Damon Runyon Cancer Research Foundation Fellowship (LVB)

Boehringer Ingelheim Fellowship (JL)

Howard Hughes Medical Institute (DCP)

National Human Genome Research Institute Intramural Research Program (MM)

NIH/NCATS Colorado CTSA grant UL1 TR002535 (NRT)

Contents are the authors’ sole responsibility and do not necessarily represent official NIH views.

Philanthropic gifts from:

Brit and Alexander d’Arbeloff

Arthur W. and Carol Tobin Brill

Matthew Brill

Charles Ellis

The Barakett Foundation

The Howard P. Colhoun Foundation

The Seedling Foundation

## Author contributions

Conceptualization: AKSR and DCP

Data curation: AKSR and HS

Formal analysis: AKSR, HS, AKG, NVB, LT, IS, and LVB

Funding acquisition: AKSR, NVB, LVB, SMD, NRT, MM, and DCP

Investigation: AKSR, TP, JL, NK, LB, SP, AD, and EP

Methodology: AKSR, HS, AKG, LT, DWB, TP, JL

Project administration: AKSR, JFH, LB, AB, PK, NB, PCL, CK, and SMD

Resources: AB, PK, NB, PCL, CK, SMD, AEL, NRT, CSS, and MM

Software: AKSR, HS, AKG, NVB, LT, IS, and LVB

Supervision: AKSR, JFH, LB, AEL, NRT, CSS, MM, and DCP

Validation: HS, AKG, LVB, DWB

Visualization: AKSR, AKG, and IS

Writing - original draft preparation: AKSR and DCP

Writing - review and editing: AKSR, HS, AKG, NVB, LT, IS, LVB, DWB, JFH, and DCP

## Inclusion and diversity

We support inclusive, diverse, and equitable conduct of research.

## STAR Methods

### RESOURCE AVAILABILITY

#### Lead contact

Further information and requests for resources and reagents should be directed to and will be fulfilled by the lead contact, David C. Page.

#### Materials availability

Cell lines generated in this study are available upon request to the lead contact.

#### Data and code availability

- Raw RNA-seq data has been deposited to dbGAP and processed data has been deposited at Github. Both are publicly available as of the date of publication. Accession numbers and DOIs are listed in the key resources table.
- This paper analyzes existing, publicly available data. Accession numbers for these datasets are listed in the key resources table.
- Original code has been deposited at Github and is publicly available as of the date of publication. The accession number is listed in the key resources table.
- Any additional information required to reanalyze the data reported in this paper is available from the lead contact upon request.

### EXPERIMENTAL MODEL AND SUBJECT DETAILS

#### Human subjects

As part of an IRB-approved study at the NIH Clinical Center (12-HG-0181) and Whitehead Institute/MIT (Protocol # 1706013503), we recruited human subjects through the NIH Clinical Center, the Focus Foundation, and Mass General Hospital. We included individuals with a previous karyotype showing non-mosaic sex chromosome aneuploidy or a structural variant of the sex chromosomes, as well as euploid “healthy volunteers”. We obtained informed consent from all study participants. For derivation of cell cultures and analysis, we collected blood samples and/or skin biopsies and shipped them to the Page lab. We performed karyotyping of peripheral blood and fibroblast cell cultures at the National Human Genome Research Institute Cytogenetics and Microscopy Core. We obtained additional cell lines from the Colorado Children’s Hospital and Coriell Research Institute, and cultured them for at least two passages prior to RNA-sequencing. The analysis presented represents the combined analysis of samples published for the first time here, and a set of previously-published samples collected during the course of the same study.^15^ We provide information about all samples analyzed in this study in **Table S1**.

### METHOD DETAILS

#### Cell culture

We derived and cultured lymphoblastoid cell lines (LCLs) and skin fibroblasts as previously described.^15^ Briefly, we collected blood in BD Vacutainer ACT blood collection tubes and placed two 4mm skin punch biopsies from the upper arm in transport media (DMEM/F12, 20% FBS, and 100 IU/ml Penicillin-Streptomycin [all ThermoFisher]) and shipped these to the Page lab 1-3 days after collection. We subjected blood to density gradient centrifugation to obtain lymphocytes, which we then transformed by culturing in complete RPMI (RPMI 1640 [Gibco], 25mM HEPES, 15% FBS [Hyclone], 1.25ug/ml Fungizone [Gibco], 3.33ug/ml Gentamycin [Gibco], 100 IU/ml Penicillin-Streptomycin [Lonza], pH 7.2) supplemented with EBV and 66.6 ug/ml cyclosporine until we observed cell clumping and proliferation. We minced skin biopsies and placed them on gelatin-coated plates with growth media (High Glucose DMEM [Gibco], 20% FBS [Hyclone], 2mM L-Glutamine [MP Biomedicals], 1X MEM Non-Essential Amino Acids [Gibco], 100 IU/ml Penicillin-Streptomycin [Lonza]), leaving undisturbed until fibroblasts grew out of the biopsies and could be passaged. For each sample, we collected and froze one million cells in RNAprotect Cell Reagent (Qiagen) at −80°C until RNA extraction. We confirmed that cell cultures were negative for mycoplasma contamination periodically using either the MycoAlert Kit (Lonza) or PCR, as described previously.^15^

#### RNA extraction and RNA-seq library preparation

We extracted RNA using the RNeasy Protect Cell Mini Kit (Qiagen), per the manufacturer’s instructions, using the following modifications: we added 10 μL β-mercaptoethanol per ml of buffer RLT for cell lysis, and 10 μL of a 1:100 dilution of ERCC RNA Spike-In Mix (Invitrogen) per million cells. We homogenized lysates using QIAshredder columns (Qiagen), removed genomic DNA using gDNA eliminator columns, performed all optional spin steps, and eluted RNA in 30 μL RNase-free water. We quantified RNA using a Qubit 2.0 or 4.0 fluorometer and the Qubit RNA HS Assay Kit (ThermoFisher). RNA was consistently high-quality with RNA integrity numbers (RIN) near 10 as measured on a Bioanalyzer 2100 instrument (Agilent). We prepared RNA sequencing libraries using the TruSeq RNA Library Preparation Kit v2 (Illumina) with modifications as detailed in Naqvi *et al.*^13^ or using the KAPA mRNA HyperPrep Kit (Roche). In both cases, we size-selected libraries using the PippinHT system (Sage Science) and 2% agarose gels with a capture window of 300-600bp. We performed paired-end 100×100 bp sequencing on a HiSeq 2500 or NovaSeq 6000 (Illumina). We indicate the library preparation kit and sequencing platform for each sample in **Table S1**.

#### RNA-seq data processing

We performed all analyses using human genome build hg38, and a custom version of the comprehensive GENCODE v24 transcriptome annotation as in Godfrey *et al*.^46^ This annotation represents the union of the “GENCODE Basic” annotation and transcripts recognized by the Consensus Coding Sequence project.^47^ To analyze samples in which ERCC spike-ins were added, we merged our custom transcript annotation with the ERCC Control annotation.

To pseudoalign reads to the transcriptome annotation and estimate expression levels of each transcript, we used kallisto software (v0.42.5).^48^ We included the --bias flag to correct for sequence bias. We imported the resulting count data (abundance.tsv file) into R for analysis in DESeq2 (v1.26.0)^16^ using the tximport package (v1.14.0).^49^ Although we pseudoaligned and performed DESeq2 on the entire custom transcriptome annotation, we restricted our downstream analysis to protein-coding and long non-coding RNA genes, as annotated in ensembl v107. We filtered the final analyses for genes with median expression level in XX or XY samples of at least 1 transcript per million (TPM) in a given cell type.

#### Linear modeling to identify autosomal X- or Y-responsive genes

We performed linear modeling in DESeq2 with three covariates: Chr X copy number, Chr Y copy number, and library preparation batch. The “log_2_ fold change” values output from DESeq2 in these analyses refer to the log_2_ fold change in expression per copy of Chr X or Y. We considered genes with P_adj_<0.05 as significantly Chr X-responsive or Chr Y-responsive.

To control for the possibility that bulk differences in sex chromosome gene expression may lead to global effects during normalization, we performed an identical analysis, removing Chr X and Y genes prior to DESeq2 normalization and linear modeling. This had little effect on the overall results.

We compared significantly X-responsive or Y-responsive genes between LCLs and fibroblasts by first restricting our analysis to autosomal genes expressed in both LCLs and fibroblasts, and then intersecting the significant gene lists. To compute P values of the overlaps, we used hypergeometric tests, with P<0.05 considered significant. We calculated Pearson correlations between log_2_ fold change per X or Y Chr in LCLs and fibroblasts on the overlapping sets of genes.

##### Saturation analysis of autosomal X- and Y-responsive genes

For LCLs and fibroblasts separately, we randomly sampled without replacement size-*n* subsets of available RNA-seq libraries. We ran DESeq2 on these subsets as in the full analysis, and recorded the number of significantly Chr X- and Y-responsive expressed autosomal genes (P_adj_<0.05). We performed 100 down-samplings for each sample size, *n*.

##### Comparing X- or Y-responsive genes to estrogen or androgen receptor target genes

We defined direct target genes of the estrogen and androgen receptors as genes that change in expression and are bound by estrogen receptor alpha (ERα) or androgen receptor (AR) within 30 kb of the transcription start site upon addition of 17-β estradiol (E2) or dihydrotestosterone (DHT), respectively. For ERα target genes, we reanalyzed RNA-seq and ChIP-seq data in MCF7 breast cancer cells from Guan *et al.*^17^, and for AR target genes we reanalyzed RNA-seq and ChIP-seq data in LNCaP prostate cancer cells from Cato *et al*.^18^ For RNA-seq data, we used the same processing pipeline as above – pseudoalignment and quantification of transcript abundance with kallisto and differential expression analysis (treatment versus vehicle) in DESeq2 – with genes with P_adj_<0.05 considered significantly responsive to E2 or DHT.

For AR and ER ChIP-seq data, we aligned the reads to the human genome (hg38) using bowtie2 (v2.3.4.1)^50^ with one mismatch allowed (-N 1) and called peaks using the MACS2 (v2.2.7.1)^51^ callpeak function with default parameters. We called peaks in hormone treated data and vehicle control, using the matched input samples as the background control. We then selected peaks exclusively found in the treated condition compared to the vehicle control. We associated peaks with genes in our custom GENCODE v24 transcriptome annotation using the bedtools (v2.26.0) closest function with default parameters. We considered genes with a peak within 30 kb of their transcription start site “bound” by a given factor.

##### Functional annotation

We performed functional gene category analysis on autosomal genes significantly responsive to Chr X or Y copy number using the Molecular Signatures Database (MSigDB) Gene Ontology Biological Process categories, version 2022.1.Hs.^52,53^ To test for enrichment of gene categories, we performed hypergeometric tests using all expressed autosomal genes in each cell type as the background set. We corrected the resulting P values for multiple hypothesis testing using the Benjamini-Hochberg method, with categories below the FDR threshold of 0.05 considered significantly enriched.

##### Chromosome 21 copy number analysis

To assess and compare the response to aneuploidy for an autosome with our sex chromosome aneuploidy model, we obtained six lymphoblastoid cell lines with Trisomy 21 from the Coriell Institute for Medical Research (AG09802, AG10316, AG10317, AG17485, AG13455, and AG16945; see **Table S1** for more details). We cultured these LCLs in the laboratory for two passages and performed RNA-sequencing in parallel with the sex chromosome aneuploidy lines. We processed raw RNA-sequencing data as above. We used data from 46,XX, 46,XY, 47,XX+21, and 47,XY+21 to evaluate the effects of three versus two copies of Chr 21. We performed this analysis in DESeq2 using three covariates in a linear model: sex chromosome constitution (XX vs XY), Chr 21 copy number (two vs three), and library preparation batch. We previously analyzed effects of Chr 21 copy number on the expression of genes encoded on Chr 21, finding that ~3/4 significantly increased in expression.^15^ Here, we compare the effects of Chr 21 copy number on the other autosomes to those of Chr X or Chr Y copy number.

#### Meta-analysis of published Xi expression datasets to define “Xa-expressed” genes

To define “Xa-expressed” genes, which have no evidence of expression from Xi, we compiled data from five studies of Chr X allelic ratios for 965 annotated protein-coding and long non-coding RNAs.^8,15, 19–21^

The first dataset (Additional file 7 in Cotton *et al*)^19^ was derived from paired genomic and cDNA SNP-chips in skewed LCL and fibroblast cell cultures. We used the provided mean AR values for 416 genes informative in at least 5 samples, their standard deviations, and the number of informative samples, to test whether the AR values were significantly greater than zero with one-sample, one-sided t-tests. We corrected the resulting P values for multiple comparisons with the p.adjust function in R using the Benjamini-Hochberg method. Genes with adjusted P values (Padj) below 0.05 were considered expressed from Xi (“escape”).

The second dataset was derived from bulk or single cell RNA-seq of LCLs (Table S5 and S8 from Tukiainen *et al*).^8^ The bulk RNA-seq was from one individual in the GTEx dataset with 100% skewed X inactivation across the body, and the single-cell RNA-seq in LCLs was from three individuals (we excluded data from one dendritic cell sample). We included 82 genes with data from at least two individuals in the single-cell dataset, or one individual in the single-cell dataset and data from the bulk RNA-seq dataset. Using the ratio of read counts from the more lowly and highly expressed alleles in each sample, we calculated AR values and used the provided adjusted P values to identify genes with significant Xi expression (Padj < 0.05). We called a gene as “escape” if one or both of the datasets showed evidence of Xi expression.

The third dataset (Dataset 3 from Garieri *et al*)^20^ was derived from single-cell allelic expression in fibroblasts.^20^ We included 203 genes that had data from at least two of the five samples in the dataset. We converted the reported values (Xa reads/total reads) to AR values using the following formula: 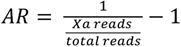. We used the provided AR threshold to consider a gene significantly expressed from Xi in each sample (AR>0.0526). We called a gene as “escape” if it had at least one sample with significant expression from Xi.

The fourth dataset (Tables S4 and S5 from Sauteraud *et al*)^21^ was derived from allele-specific bulk RNA-seq performed on 136 LCLs with skewed X inactivation. We analyzed 215 genes that had data from at least 10 samples. We calculated an AR value for each gene in each sample using the read counts from the more lowly and highly expressed alleles in each sample, adjusting for the level of skewing in each sample. To identify genes that were significantly expressed from Xi across samples, we performed paired, two-sample, one-sided t-tests, asking whether the raw (pre-adjusted for skewing) AR values were greater than the baseline AR given the level of skewing in each sample 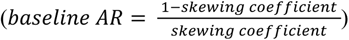; we corrected the resulting P values for multiple comparisons with the p.adjust function in R using the Benjamini-Hochberg method. We considered genes with Padj < 0.01 as significantly expressed from Xi (“escape”).

The fifth dataset (Table S6 from San Roman *et al*)^15^ was derived from our prior analysis of allelic ratios in bulk RNA-sequencing of skewed LCLs and fibroblasts with two copies of the X chromosome. We calculated AR values for 152 genes in LCLs and 120 in fibroblasts. We used the Benjamini-Hochberg adjusted P values from one-sided, one-sample t-tests to determine whether AR values were significantly greater than zero, considering genes with Padj < 0.05 as expressed from Xi (“escape”).

Next, we synthesized the calls from these five datasets. We assigned a gene as having evidence for expression from Xi (“escape”) if 1) most (>50% of) studies indicated “escape” or 2) 50% or fewer (but more than 0) studies indicated “escape” and either i) there was more than one study with evidence of escape or ii) the average AR across all studies was ≥0.1. We assigned genes as not expressed from Xi (“subject”) if: 1) all studies indicated “subject” or 2) most (>50%) studies indicated “subject” and the average AR across all studies was <0.1. For genes with no data from any of the five studies, we investigated whether there was evidence of expression from Xi using human-rodent hybrid cell lines carrying a human Xi from Carrel *et al*^7^ as compiled in Balaton *et al* ^22^. If a gene was expressed in at least 22% of Xi hybrid cell lines (per Carrel *et al*), we considered the gene to be expressed from Xi (“escape”). Our final annotations of “Xa-expressed” genes are listed in **Table S4**. In total, we found 107 genes with prior evidence of expression from Xi (“escape”), 460 genes without prior evidence of expression from Xi (“subject” – considered here as “Xa-expressed”), and 398 genes without data to make a call (“no call”).

##### Analysis of Yq heterochromatin length

To assess Yq heterochromatin length, we analyzed whole genome sequencing data of 1,225 males from the 1000 Genomes Project.^25^ We aligned FASTQ files with paired-end reads to the human reference genome (hg38) using bowtie2.^50^ From each male in the sample, we calculated the average depth of coverage of the *DYZ1* region. In addition, we calculated the average depth of coverage of a 1-megabase single-copy region of the Y chromosome (from bases 14,500,000-15,500,000). We adjusted read depths to account for GC bias in sequencing depth.^54,55^ In cases of individuals with multiple sequenced samples, we summed the read depths following GC-bias correction. For each male, we normalized mean *DYZ1* read depth by mean single-copy region read depth to account for differences in sequencing depth between different sequencing runs. After these steps, a higher depth of coverage implies higher *DYZ1* copy number and an increase in the size of the Yq heterochromatic region. We confirmed these size differences visually using FISH (see below) on eight males from the 1000 Genomes Project (NA20520, HG04185, HG01890, NA12812, HG02684, NA18748, HG02394, and HG00142) obtained from the NHGRI Sample Repository for Human Genetic Research at the Coriell Institute for Medical Research.

To analyze gene expression as a function of *DYZ1* read depth, we obtained RNA-seq data from The Geuvadis Consortium and restricted our analysis to samples marked as male, “UseThisDuplicate == 1”, RNA integrity number (RIN) ≥ 8, and with estimates for *DYZ1* read depth. We excluded the following outlier or incorrectly annotated samples as flagged in t’Hoen *et al*.^56^: NA12546.1, HG00329.5, NA18861.4, NA19144.4, NA19225.6, HG00237.4, NA12399.7, and NA07000.1, resulting in a total of 194 male samples. We downloaded the raw RNA-seq data and pseudoaligned the reads with kallisto as above. Using DESeq2, we modeled the log_2_ read counts as a function of log_2_ *DYZ1* depth of coverage, controlling for population and the lab that sequenced the libraries.

##### Analysis of gene expression in individuals with structural variants of the sex chromosomes

Naturally occurring structural variants of the sex chromosomes allowed us to investigate the consequences of deleting or duplicating specific genes on Chr X or Chr Y. The Page lab collected these samples over several decades during the course of studying infertility, disorders of sex development, and sex chromosome aneuploidy. We cultured these cells and performed RNA-sequencing in parallel with the sex chromosome aneuploidy cell lines.

##### Y isochromosomes

The Page lab previously identified three unrelated individuals with Y isochromosomes lacking the Yq heterochromatic region among males with spermatogenic failure.^57,58^ One of these isochromosomes resulted from recombination between the IR4 repeats on Yp and Yq (WHT5557, referred to as a “pseudoisochromosome”) and two resulted from recombination more distally, within the P1 palindrome. Both recombination events resulted in the duplication of protein-coding NPY genes and all PAR1 genes expressed in somatic cell types. For the 442 X- and Y-responsive autosomal genes in LCLs, we used PCA plots to visualize the clustering of the Y isochromosome samples within the Chr Y chromosome copy number series, controlling for a single X chromosome (0Y: 45,X; 1Y: 46,XY; 2Y: 47,XYY; 4Y: 49,XYYYY). For the PCA plots, we performed a variance stabilizing transformation on normalized counts from DESeq2 and used the removeBatchEffect function in the limma R package (v3.42.2)^59^ to remove any effects of library preparation batch. Ellipses representing the 95% confidence intervals around the centroid of each group (with at least 3 samples) were computed using ClustVis software.^60^

##### X isochromosomes

Through karyotyping of individuals with Turner syndrome, we identified four unrelated individuals with X isochromosomes in which the entirety of Xq – and in some cases proximal portions of Xp – is duplicated. We mapped the breakpoints of these isochromosomes by fluorescent in situ hybridization (FISH) using bacterial artificial chromosomes (BACs) with known locations across the X chromosome as DNA probes (**Fig. S11**; see below for FISH methods). In two individuals, WHT7183 and WHT7399, the breakpoints mapped to the centromere. In these two cases, probe 343B12, proximal to all genes on Xp, was absent on the isochromosome, and probe 168B01, proximal to all genes on Xq, was present in two copies. A third individual, WHT7182, had a breakpoint on Xp ~5 Mb from the centromere. This breakpoint mapped within the 206-kb region (chrX:52,607,059-52,813,686) covered by FISH probe 34G18. This region contains a palindrome (P5), the likely site of homologous recombination. A fourth individual, WHT7207, had a breakpoint on distal Xp near PAR1. We mapped the breakpoint to a 361-kb region (chrX:9,438,772-9,842,745) from *TBL1X* to *SHROOM2. TBL1X* is expressed at a level consistent with it being present on this individual’s isochromosome, indicating that the breakpoint is proximal to *TBL1X*.

In analyzing gene expression from the X isochromosome samples, we discovered that the isochromosome was not always stable in the cell cultures. Samples with recombination in the centromere had stable karyotypes in both LCLs and fibroblasts. However, for the two samples with more distal Xp breakpoints, LCLs mostly or completely lost the isochromosome and thus could not be used in this analysis. Fibroblasts retained the isochromosome in enough cells that we could analyze their gene expression (WHT7182: 47% 46,X,i(Xq) and 53% 45,X, 49 interphase nuclei; WHT7207: 64% 46,X,i(Xq) and 36% 45,X, 100 interphase nuclei;), however the resulting expression levels should be considered an average between these populations. For the X- and Y-responsive autosomal genes in LCLs or fibroblasts, we used PCA plots to visualize the clustering of the X isochromosome samples with the Chr X copy number series, without Y chromosomes (1X: 45,X; 2X: 46,XX; 3X: 47,XXX; 4X: 48,XXXX).

##### X-Y translocations

Among individuals with 46,XY disorders of sexual development or Turner syndrome, we identified three females (WHT1297, WHT1489, WHT1615) with X-Y translocations between the *PRKX* and *PRKY* genes [46,X,t(X;Y)(*PRKX-PRKY*)]. Recombination between *PRKX* and *PRKY* is common among 46,XY females and is facilitated by a polymorphic Yp inversion.^24,61^ We used a PCR-based screening approach to determine and sequence the breakpoints (**Table S7**). These females are missing five NPX-NPY genes: *SRY, RPS4Y1, ZFY, AMELY, and TBL1Y.* Samples WHT1297 and WHT1489 are missing *PRKX* but retain *PRKY*, while sample WHT1615 is missing *PRKY* but retains *PRKX*.

Also among individuals with 46,XY disorders of sexual development, we ascertained four females (WHT1785, WHT1798, WHT1799, WHT2315) with X-Y translocations between the *ANOS1 (KAL1)* and *ANOS2P (KALP)* genes [46,X,t(X;Y)(*ANOS1-ANOS2P*)]. We identified breakpoints using a PCR-based screening approach (**Table S7**). In three of the cases (WHT1785, WHT1798, and WHT1799, *ANOS1* is retained, while *ANOS2P* is deleted. In WHT2315, the recombination is within *ANOS1* and *ANOS2P*, resulting in the partial deletion of each. Based on RNA-seq data of the translocated samples, some or all of the of the genes on Yq are not expressed. This is likely due to spreading of X chromosome inactivation onto the translocated portion of the Y. In WHT1799, no Y genes are expressed; in WHT1785 and WHT1798 only *EIF1AY* is expressed; while in WHT2315 *TXLNGY* and *KDM5D* are expressed at low levels and *EIF1AY* is expressed (**Table S7**).

For the X- and Y-responsive autosomal genes in LCLs, we used PCA plots to visualize the clustering of the X-Y translocation samples with the Chr X and Y copy number series (1X: 45,X; 2X: 46,XX; 2X: 47,XXX; 4X: 48,XXXX; 1Y: 46,XY; 2Y: 47,XYY; 4Y: 49,XYYYY).

##### Fluorescent in situ hybridization

To assess copy number and variations in X and Y chromosome structure, we performed DNA fluorescent *in situ* hybridization using BACs with known locations as DNA probes, as previously described.^57^ We labelled FISH probes by nick translation with biotin-dUTP (Sigma) and Cy3-dUTP (Amersham). To detect Biotin-labelled probes, we used Avidin-FITC probes (Life Technologies, 1:250 dilution), and counterstained DNA with DAPI. We captured images on a DeltaVision deconvolution microscope. We provide information on probes for FISH experiments in **Table S7**. For copy number analysis, we performed FISH on interphase cells using Whole X and Whole Y probes (Applied Spectral Imaging).

##### Motif analysis

We analyzed DNA motif enrichment using Homer software v4.11.1.^62^ The findMotifs function identified *de novo* motifs enriched in promoters of a gene set of interest (e.g. Chr X-responsive autosomal genes in LCLs) relative to expressed genes as a “background” set (e.g., autosomal genes expressed in LCLs). We defined promoters as +/- 1 kb from the transcription start site. We used the “-fdr” argument with 100 randomizations to calculate an empirical FDR for the motif enrichments. We then compared the enriched motifs to motif databases to identify the best match among known DNA binding proteins.

#### Analysis of ZFX ChIP-seq data

To detect ZFX binding at gene promoters, we analyzed four ZFX ChIP-seq experiments in different cell lines from ENCODE4 v1.6.1 (HCT 116, C4-2B, MCF-7, and HEK-293T). Using the BEDTools closest function, we mapped ZFX peaks to the nearest annotated transcription start site in the GENCODE v24 comprehensive annotation.^63^ For gene expression analysis in each cell line, we processed fastq files from ENCODE or GEO^29,36^ using kallisto as above or, when available, we downloaded pre-processed RNA-seq data. We restricted our analysis of ZFX peaks in each cell line to genes with median TPM≥1.

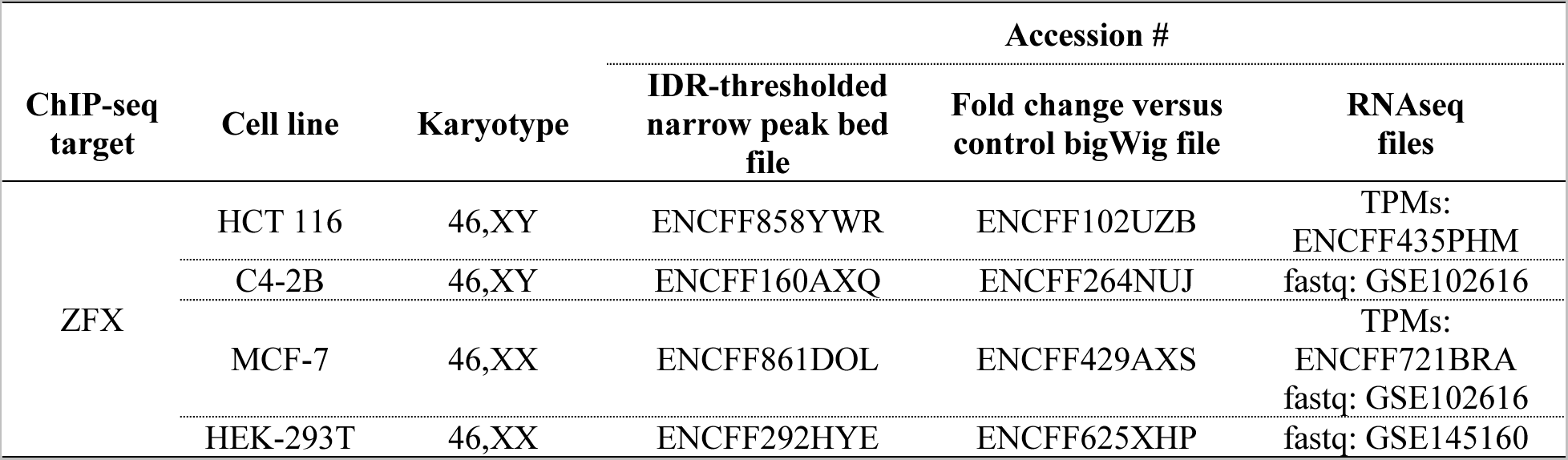

To evaluate ZFX binding in promoters of genes, we investigated whether there was a ChIP-seq peak 1 kb up- or downstream of the transcription start site. To get a sense for the most robust binding sites across cell types, we calculated the proportion of cell lines bound at the promoter of each gene by dividing the number of cell lines with binding by the number of cell lines in which that gene is expressed.

#### Generating cell lines for CRISPR interference

To knock down endogenous *ZFX* or *ZFY* levels in 46,XX and 46,XY fibroblasts, we used CRISPR interference to target a nuclease-dead Cas9 fused with a repressive KRAB domain (dCas9-KRAB) to the *ZFX* or *ZFY* promoter and thereby repress *ZFX* or *ZFY* transcription.

##### Lentiviral production

We obtained the following constructs from Addgene for use in our experiments: 1) for CRISPRi, pLX_311-KRAB-dCas9 (Addgene: #96918), a KRAB-dCas9 blasticidin-selectable lentiviral expression vector;^37^ 2) sgOpti (#85681), a puromycin-selectable structurally optimized guide RNA (gRNA) lentiviral expression vector;^64,65^ 3) psPAX2 (#12260), a second generation lentiviral packaging plasmid; and 4) pCMV-VSV-G (#8454), a lentiviral envelope protein plasmid.^66^ We purified all plasmids using the EndoFree Maxi Kit (Qiagen).

We used the top five guide sequences for *ZFX* and *ZFY* from the human CRISPRi v2 gRNA library.^67^ We tested these guides for *ZFX* or *ZFY* knockdown and chose the two guides that gave the most robust response for the final experiments. We also used intragenic (IG) control guides mapping to a gene-poor region on Chr 2.^68^ Guides and primer sequences are listed below.

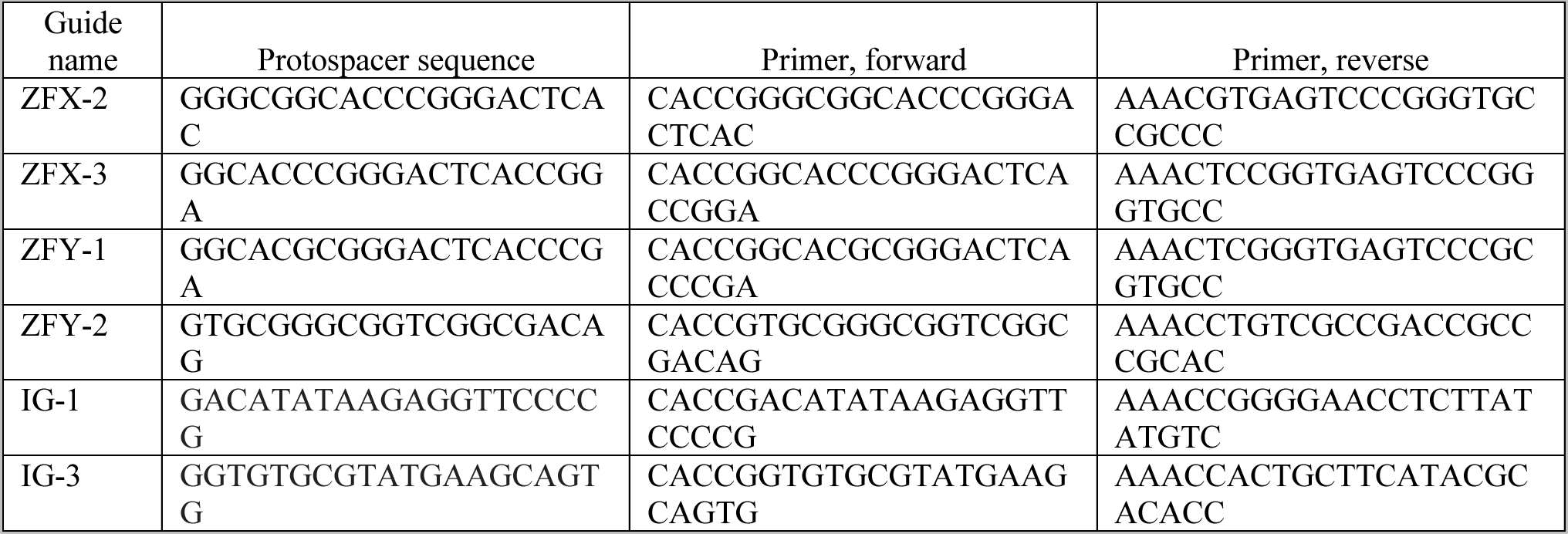

To clone guides into the sgOpti vector, we digested using FastDigest BsmBI (ThermoFisher) and dephosphorylated the ends with FastAP (ThermoFisher) for 30 min at 37°C. We gel-purified the digested plasmid using the QIAquick Gel Extraction Kit (Qiagen). Prior to ligation, we phosphorylated and annealed each pair of oligos using T4 Polynucleotide kinase (New England BioLabs). We then ligated each insert into the backbone using Quick Ligase (New England BioLabs) for 10 min at room temperature, and transformed into NEB Stable Cells for amplification (New England BioLabs). We confirmed gRNA sequences by sequencing.

LentiX-293T cells (Takara) were cultured in DMEM with 2mM L-Glutamine, 100U/mL Penicillin-Streptomycin, and 10% FBS on plates coated with collagen (5 μg/cm^2^) to produce virus. For gRNA constructs, we plated 5×10^6^ LentiX-293T cells in one 10 cm plate the night before transfection. The next day, we co-transfected 4.5 μg of sgOpti, 4 μg of pCMV-VSV-G, and 6.5 μg of psPAX2 using TransIT-LT1 reagent (Mirus). We harvested virus-containing media once at 48 h and once at 72 h, pooled, and tested for successful viral production using Lenti-X GoStiX (Takara). We concentrated gRNA virus 10X using Lenti-X concentrator (Takara). For the dCas9-KRAB and dCas9-VP64 constructs, we co-transfected five plates of LentiX-293T cells with 7 μg of pLX_311 or pXPR_120, 3 μg of pCMV-VSV-G, and 5 μg of psPAX2. We added Viral Boost reagent (Takara) 24 h post transfection, and again with replacement media after harvesting at 48 h. We pooled virus-containing media across all plates and concentrated 100X.

##### Generation of dCas9-KRAB and dCas9-VP64 fibroblast cultures

For CRISPRi we transduced 46,XX and 46,XY fibroblasts from three individuals each. We performed transductions using the “spinfection” method: we added 1 mL fibroblast media with 5-20 uL of 100X concentrated pLX_311 lentiviral vector and 8 μg/mL polybrene (hexamethrine bromide; Sigma) to 7.5×10^4^ cells per well in 12-well plates or 1.875×10^5^ cells per well in 6-well plates, spun at 1000 rpm for 1 h and incubated overnight. 24 h post infection, we trypsinized each and expanded to 6-well or 10-cm plates with fibroblast media containing 5 μg/mL blasticidin (Life Technologies), which we maintained throughout the experiment for continuous selection.

##### *ZFX* and *ZFY* knockdown and analysis

We transduced control (intragenic) and *ZFX-* or *ZFY*-targeting gRNAs into the stably-expressing dCas9-KRAB fibroblasts as above, and selected cells using 2 μg/mL puromycin (Sigma) beginning 24 h post infection. We collected cells 72 h post infection. We isolated RNA and prepared libraries for RNA-seq as described above and sequenced using 150 bp paired-end reads on a NovaSeq 6000. We trimmed reads to 100 bp, pseudoaligned with kallisto, and imported the count data into R using tximport, as described above. We used DESeq2 to perform differential expression analysis including sgRNA target (*ZFX*, *ZFY*, or intergenic control) and cell line as covariates.

### QUANTIFICATION AND STATISTICAL ANALYSIS

We used various statistical tests to calculate P values as indicated in the methods section, figure legend, or text, where appropriate. To calculate all statistics and generate plots, we used R software, version 3.6.3.^69^ We considered results statistically significant when P<0.05 or, when using multiple hypothesis correction, P_adj_<0.05 or FDR<0.05. Unless otherwise stated, we plotted data summaries as median and interquartile range. We used Deming regressions to compare variables that measured with error (e.g., log_2_ fold change per chrX vs log_2_ fold change per chrY). We calculated Deming regressions using the R package “deming” v1.4. For weighted Deming regressions, we included error values using the “xstd” and “ystd” arguments.

### KEY RESOURCES TABLE

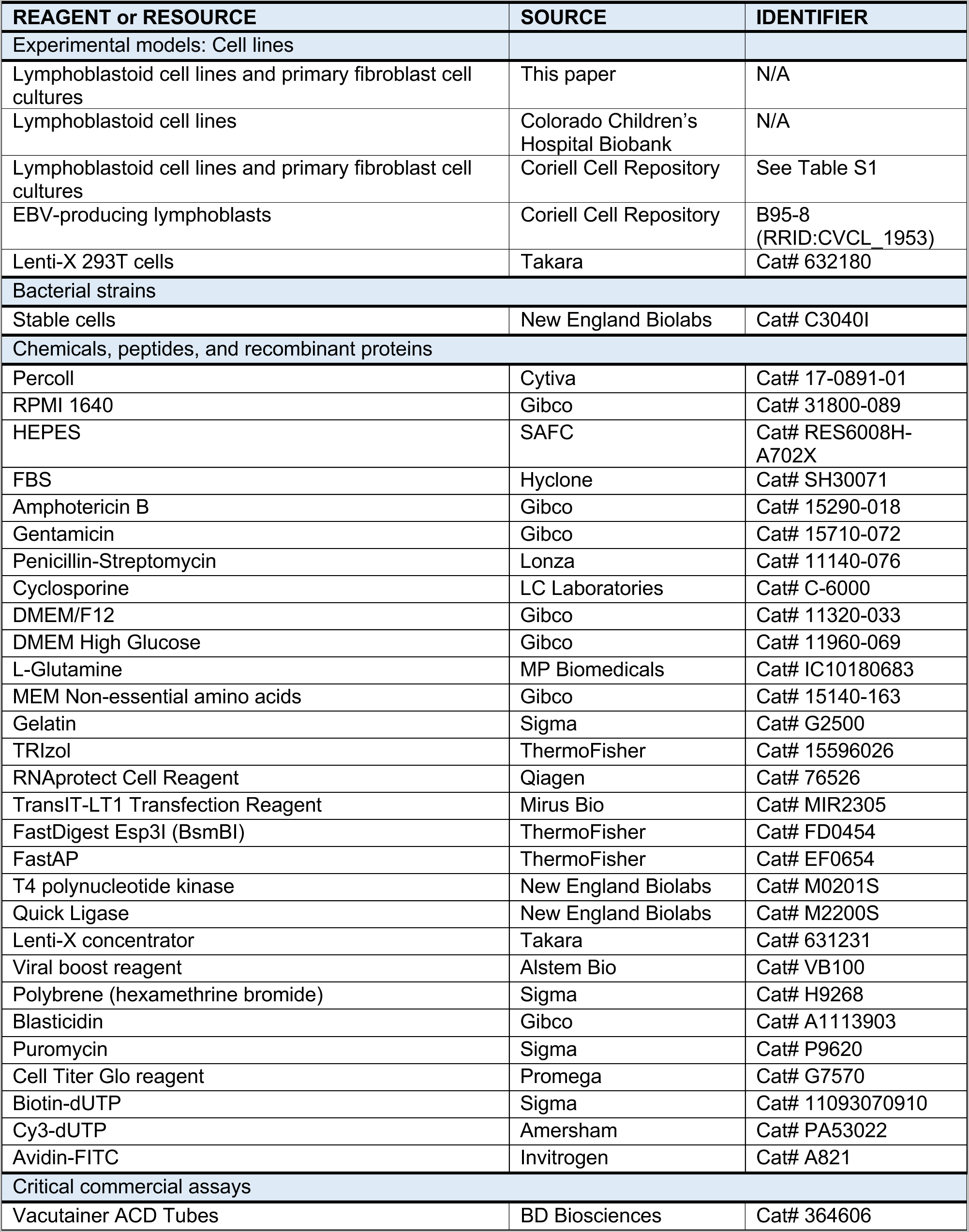
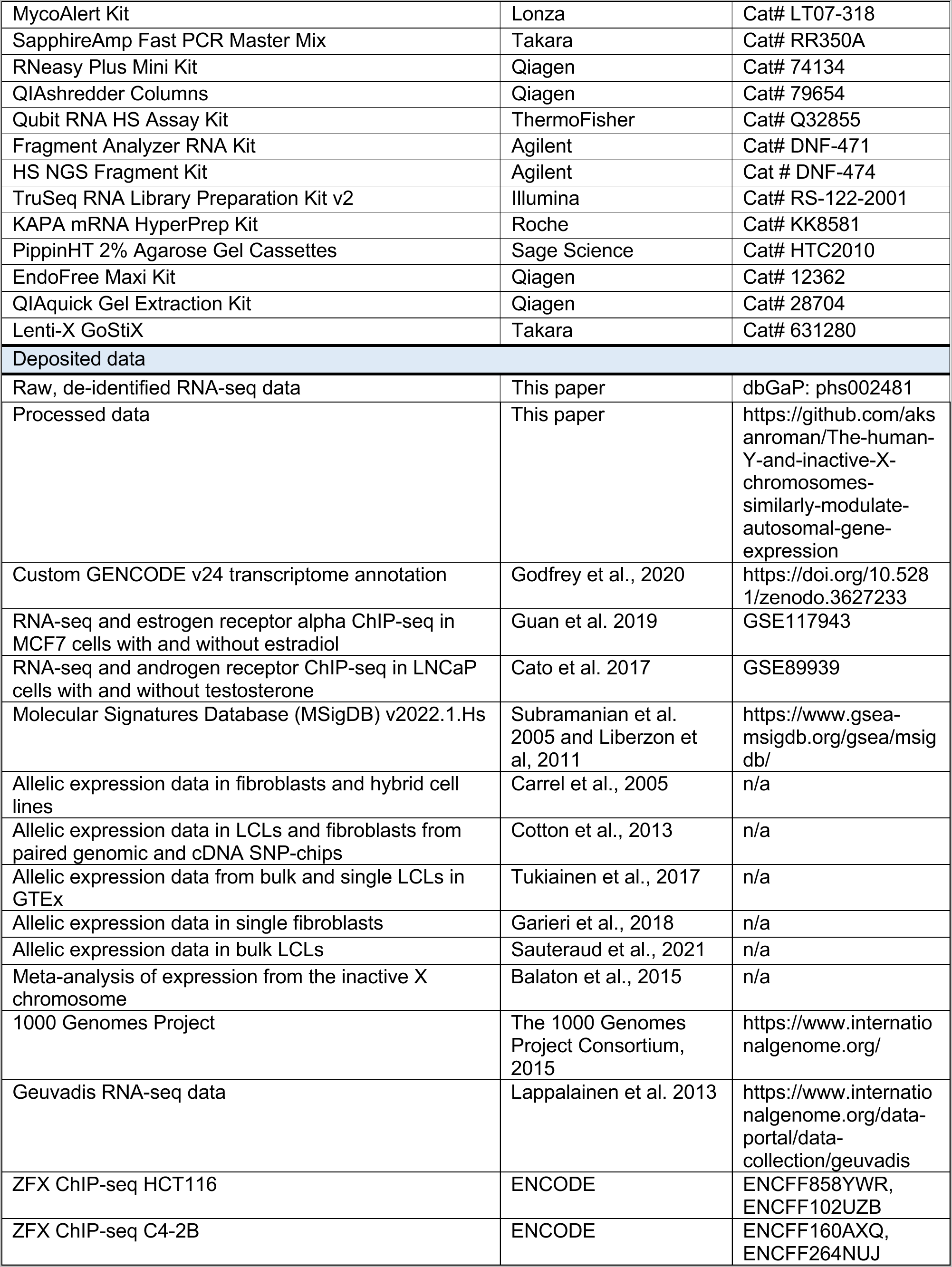
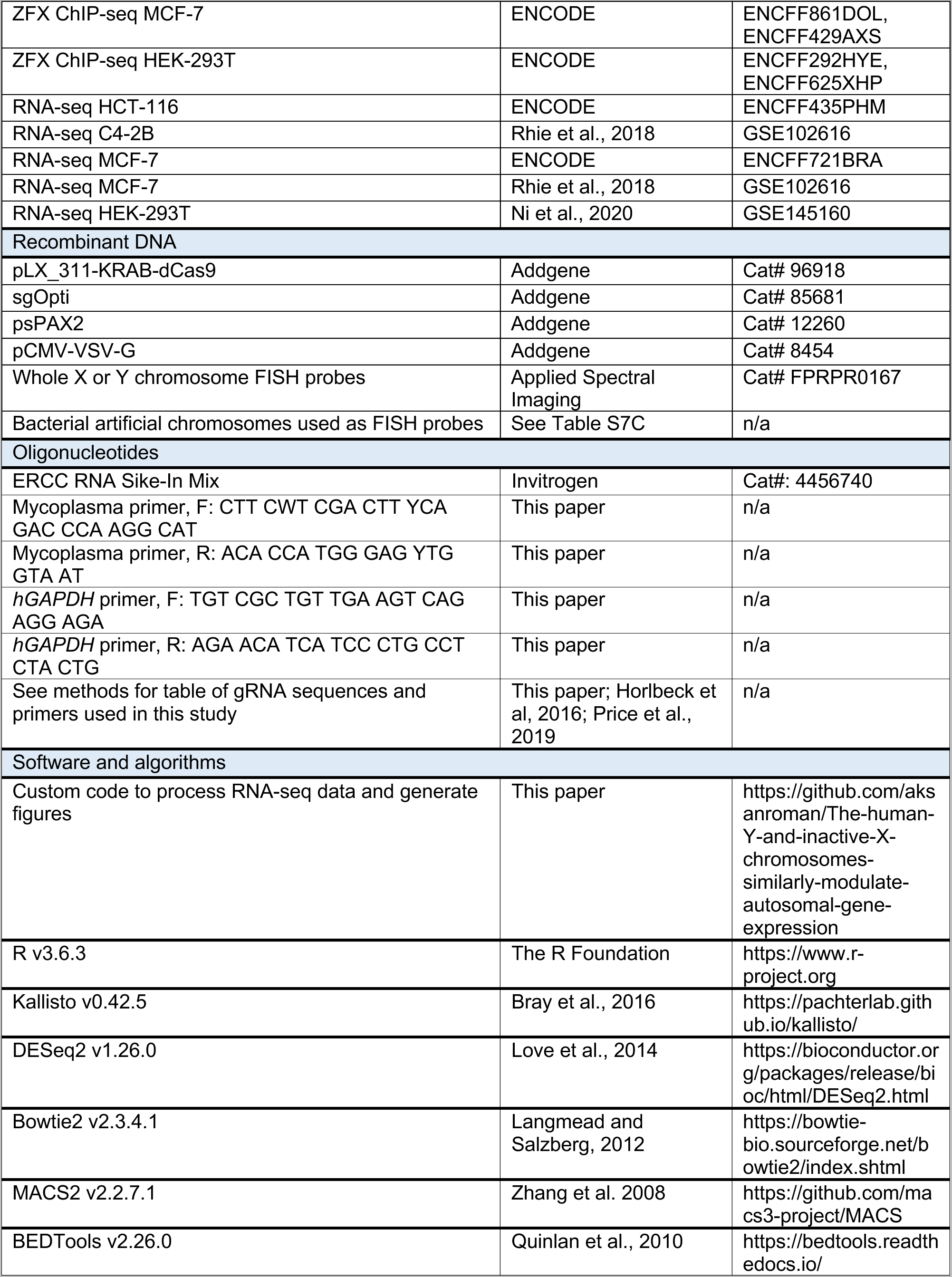
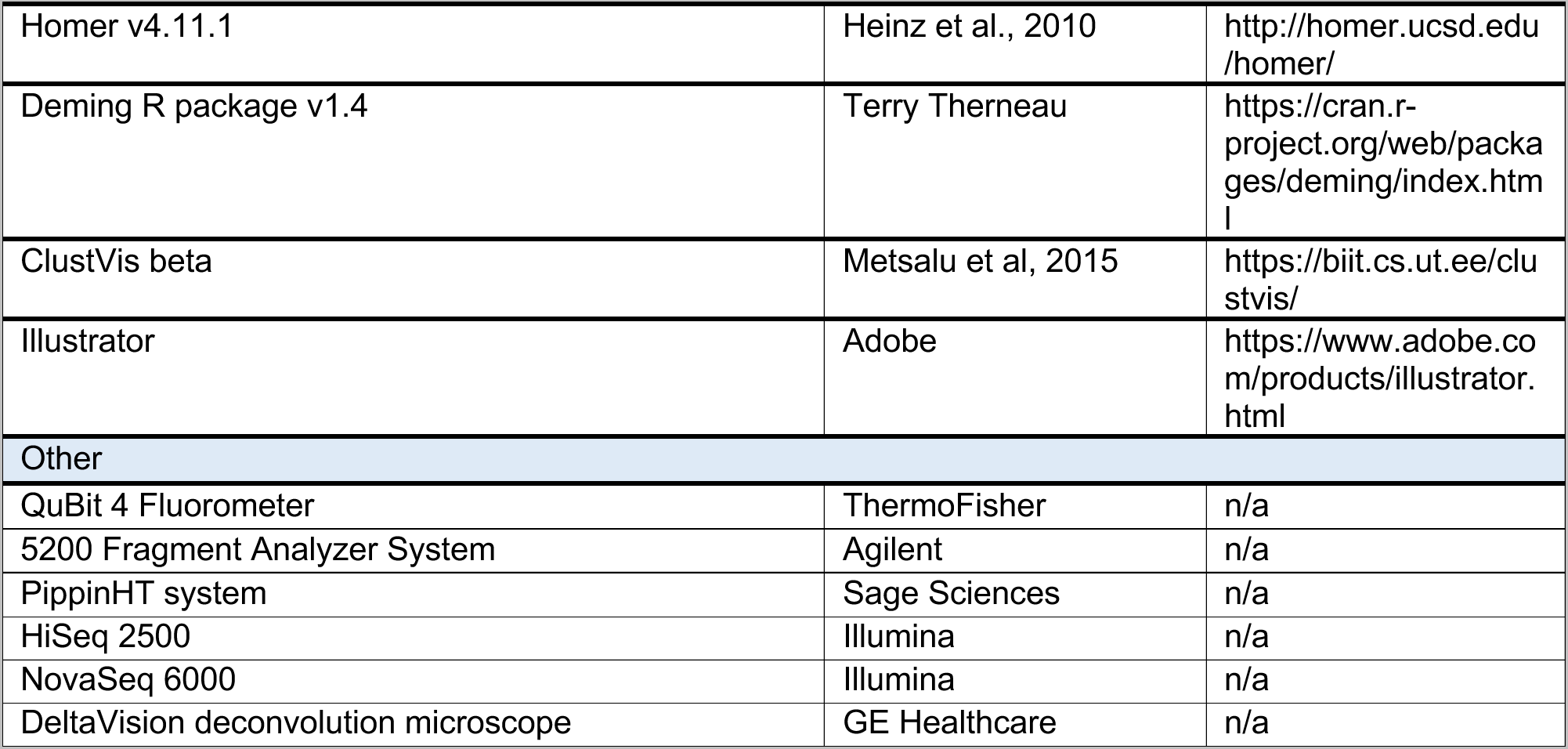

## References

1. Ohno, S. (1967). Sex Chromosomes and Sex-Linked Genes (Springer Berlin Heidelberg). 10.1007/978-3-642-88178-7.

2. Lahn, B.T., and Page, D.C. (1999). Four evolutionary strata on the human X chromosome. Science 286, 964–967. DOI 10.1126/science.286.5441.964.

3. Lyon, M.F. (1961). Gene action in the X-chromosome of the mouse (Mus musculus L.). Nature 190, 372–373. 10.1038/190372a0.

4. Brown, C.J., Ballabio, A., Rupert, J.L., Lafreniere, R.G., Grompe, M., Tonlorenzi, R., and Willard, H.F. (1991). A gene from the region of the human X inactivation centre is expressed exclusively from the inactive X chromosome. Nature 349, 38–44. 10.1038/349038a0.

5. Stern, C. (1957). The problem of complete Y-linkage in man. Am. J. Hum. Genet. 9, 147–166.

6. Skaletsky, H., Kuroda-Kawaguchi, T., Minx, P.J., Cordum, H.S., Hillier, L., Brown, L.G., Repping, S., Pyntikova, T., Ali, J., Bieri, T., et al. (2003). The male-specific region of the human Y chromosome is a mosaic of discrete sequence classes. Nature 423, 825–837. 10.1038/nature01722.

7. Carrel, L., and Willard, H.F. (2005). X-inactivation profile reveals extensive variability in X-linked gene expression in females. Nature 434, 400–404. 10.1038/nature03479.

8. Tukiainen, T., Villani, A.C., Yen, A., Rivas, M.A., Marshall, J.L., Satija, R., Aguirre, M., Gauthier, L., Fleharty, M., Kirby, A., et al. (2017). Landscape of X chromosome inactivation across human tissues. Nature 550, 244–248. 10.1038/nature24265.

9. Lahn, B.T., and Page, D.C. (1997). Functional coherence of the human Y chromosome. Science 278, 675–680. DOI 10.1126/science.278.5338.675.

10. Bellott, D.W., Hughes, J.F., Skaletsky, H., Brown, L.G., Pyntikova, T., Cho, T.-J., Koutseva, N., Zaghlul, S., Graves, T., Rock, S., et al. (2014). Mammalian Y chromosomes retain widely expressed dosage-sensitive regulators. Nature 508, 494–499. 10.1038/nature13206.

11. Forsberg, L.A. (2017). Loss of chromosome Y (LOY) in blood cells is associated with increased risk for disease and mortality in aging men. Hum. Genet. 136, 657–663. 10.1007/s00439-017-1799-2.

12. Melé, M., Ferreira, P.G., Reverter, F., DeLuca, D.S., Monlong, J., Sammeth, M., Young, T.R., Goldmann, J.M., Pervouchine, D.D., Sullivan, T.J., et al. (2015). The human transcriptome across tissues and individuals. Science (New York, NY) 348, 660–665. 10.1126/science.aaa0355.

13. Naqvi, S., Godfrey, A.K., Hughes, J.F., Goodheart, M.L., Mitchell, R.N., and Page, D.C. (2019). Conservation, acquisition, and functional impact of sex-biased gene expression in mammals. Science 365. 10.1126/science.aaw7317.

14. Oliva, M., Munoz-Aguirre, M., Kim-Hellmuth, S., Wucher, V., Gewirtz, A.D.H., Cotter, D.J., Parsana, P., Kasela, S., Balliu, B., Vinuela, A., et al. (2020). The impact of sex on gene expression across human tissues. Science 369. 10.1126/science.aba3066.

15. San Roman, A.K., Godfrey, A.K., Skaletsky, H., Bellott, D.W., Groff, A.F., Harris, H.L., Blanton, L.V., Hughes, J.F., Brown, L., Phou, S., et al. (2023). The human inactive X chromosome modulates expression of the active X chromosome. Cell Genomics 3, 100259. 10.1016/j.xgen.2023.100259.

16. Love, M.I., Huber, W., and Anders, S. (2014). Moderated estimation of fold change and dispersion for RNA-seq data with DESeq2. Genome Biol. 15, 550. 10.1186/s13059-014-0550-8.

17. Guan, J., Zhou, W., Hafner, M., Blake, R.A., Chalouni, C., Chen, I.P., De Bruyn, T., Giltnane, J.M., Hartman, S.J., Heidersbach, A., et al. (2019). Therapeutic Ligands Antagonize Estrogen Receptor Function by Impairing Its Mobility. Cell 178, 949–963 e918. 10.1016/j.cell.2019.06.026.

18. Cato, L., Neeb, A., Sharp, A., Buzon, V., Ficarro, S.B., Yang, L., Muhle-Goll, C., Kuznik, N.C., Riisnaes, R., Nava Rodrigues, D., et al. (2017). Development of Bag-1L as a therapeutic target in androgen receptor-dependent prostate cancer. Elife 6. 10.7554/eLife.27159.

19. Cotton, A.M., Ge, B., Light, N., Adoue, V., Pastinen, T., and Brown, C.J. (2013). Analysis of expressed SNPs identifies variable extents of expression from the human inactive X chromosome. Genome Biol. 14, R122. 10.1186/gb-2013-14-11-r122.

20. Garieri, M., Stamoulis, G., Blanc, X., Falconnet, E., Ribaux, P., Borel, C., Santoni, F., and Antonarakis, S.E. (2018). Extensive cellular heterogeneity of X inactivation revealed by single-cell allele-specific expression in human fibroblasts. Proc. Natl. Acad. Sci. USA 115, 13015–13020. 10.1073/pnas.1806811115.

21. Sauteraud, R., Stahl, J.M., James, J., Englebright, M., Chen, F., Zhan, X., Carrel, L., and Liu, D.J. (2021). Inferring genes that escape X-Chromosome inactivation reveals important contribution of variable escape genes to sex-biased diseases. Genome Res. 31, 1629–1637. 10.1101/gr.275677.121.

22. Balaton, B.P., Cotton, A.M., and Brown, C.J. (2015). Derivation of consensus inactivation status for X-linked genes from genome-wide studies. Biol. Sex Differ. 6. 10.1186/s13293-015-0053-7.

23. Lemos, B., Branco, A.T., and Hartl, D.L. (2010). Epigenetic effects of polymorphic Y chromosomes modulate chromatin components, immune response, and sexual conflict. Proceedings of the National Academy of Sciences of the United States of America 107, 15826–15831. 10.1073/pnas.1010383107.

24. Repping, S., van Daalen, S.K.M., Brown, L.G., Korver, C.M., Lange, J., Marszalek, J.D., Pyntikova, T., van der Veen, F., Skaletsky, H., Page, D.C., and Rozen, S. (2006). High mutation rates have driven extensive structural polymorphism among human Y chromosomes. Nat. Genet. 38, 463–467. 10.1038/ng1754.

25. The 1000 Genomes Project Consortium (2015). A global reference for human genetic variation. Nature 526, 68–74. 10.1038/nature15393.

26. Lappalainen, T., Sammeth, M., Friedländer, M.R., t Hoen, P.A.C., Monlong, J., Rivas, M.A., Gonzàlez-Porta, M., Kurbatova, N., Griebel, T., Ferreira, P.G., et al. (2013). Transcriptome and genome sequencing uncovers functional variation in humans. Nature 501, 506–511. 10.1038/nature12531.

27. Weil, D., Wang, I., Dietrich, A., Poustka, A., Weissenbach, J., and Petit, C. (1994). Highly homologous loci on the X and Y chromosomes are hot-spots for ectopic recombinations leading to XX maleness. Nat. Genet. 7, 414–419. 10.1038/ng0794-414.

28. Decarpentrie, F., Vernet, N., Mahadevaiah, S.K., Longepied, G., Streichemberger, E., Aknin-Seifer, I., Ojarikre, O.A., Burgoyne, P.S., Metzler-Guillemain, C., and Mitchell, M.J. (2012). Human and mouse ZFY genes produce a conserved testis-specific transcript encoding a zinc finger protein with a short acidic domain and modified transactivation potential. Hum. Mol. Genet. 21, 2631–2645. 10.1093/hmg/dds088.

29. Rhie, S.K., Yao, L., Luo, Z., Witt, H., Schreiner, S., Guo, Y., Perez, A.A., and Farnham, P.J. (2018). ZFX acts as a transcriptional activator in multiple types of human tumors by binding downstream from transcription start sites at the majority of CpG island promoters. Genome Res. 28, 310–320. 10.1101/gr.228809.117.

30. Mardon, G., Luoh, S.W., Simpson, E.M., Gill, G., Brown, L.G., and Page, D.C. (1990). Mouse Zfx protein is similar to Zfy-2: each contains an acidic activating domain and 13 zinc fingers. Mol. Cell. Biol. 10, 681–688.

31. Xu, S., Duan, P., Li, J., Senkowski, T., Guo, F., Chen, H., Romero, A., Cui, Y., Liu, J., and Jiang, S.W. (2016). Zinc Finger and X-Linked Factor (ZFX) Binds to Human SET Transcript 2 Promoter and Transactivates SET Expression. Int J Mol Sci 17. 10.3390/ijms17101737.

32. Raznahan, A., Parikshak, N.N., Chandran, V., Blumenthal, J.D., Clasen, L.S., Alexander-Bloch, A.F., Zinn, A.R., Wangsa, D., Wise, J., Murphy, D.G.M., et al. (2018). Sex-chromosome dosage effects on gene expression in humans. Proc. Natl. Acad. Sci. USA 115, 7398–7403. 10.1073/pnas.1802889115.

33. Zhang, X., Hong, D., Ma, S., Ward, T., Ho, M., Pattni, R., Duren, Z., Stankov, A., Bade Shrestha, S., Hallmayer, J., et al. (2020). Integrated functional genomic analyses of Klinefelter and Turner syndromes reveal global network effects of altered X chromosome dosage. Proc. Natl. Acad. Sci. USA 117, 4864–4873. 10.1073/pnas.1910003117.

34. Schneider-G dicke, A., Beer-Romero, P., Brown, L.G., Nussbaum, R., and Page, D.C. (1989). ZFX has a gene structure similar to ZFY, the putative human sex determinant, and escapes X inactivation. Cell 57, 1247–1258. 10.1016/0092-8674(89)90061-5.

35. Palmer, M.S., Berta, P., Sinclair, A.H., Pym, B., and Goodfellow, P.N. (1990). Comparison of human ZFY and ZFX transcripts. Proceedings of the National Academy of Sciences 87, 1681–1685.

36. Ni, W., Perez, A.A., Schreiner, S., Nicolet, C.M., and Farnham, P.J. (2020). Characterization of the ZFX family of transcription factors that bind downstream of the start site of CpG island promoters. Nucleic Acids Res. 48, 5986–6000. 10.1093/nar/gkaa384.

37. Rosenbluh, J., Xu, H., Harrington, W., Gill, S., Wang, X., Vazquez, F., Root, D.E., Tsherniak, A., and Hahn, W.C. (2017). Complementary information derived from CRISPR Cas9 mediated gene deletion and suppression. Nat. Commun. 8, ncomms15403. 10.1038/ncomms15403.

38. Gershoni, M., and Pietrokovski, S. (2017). The landscape of sex-differential transcriptome and its consequent selection in human adults. BMC Biol. 15, 7. 10.1186/s12915-017-0352-z.

39. Lopes-Ramos, C.M., Chen, C.Y., Kuijjer, M.L., Paulson, J.N., Sonawane, A.R., Fagny, M., Platig, J., Glass, K., Quackenbush, J., and DeMeo, D.L. (2020). Sex Differences in Gene Expression and Regulatory Networks across 29 Human Tissues. Cell Rep 31, 107795. 10.1016/j.celrep.2020.107795.

40. Trolle, C., Nielsen, M.M., Skakkebæk, A., Lamy, P., Vang, S., Hedegaard, J., Nordentoft, I., Ørntoft, T.F., Pedersen, J.S., and Gravholt, C.H. (2016). Widespread DNA hypomethylation and differential gene expression in Turner syndrome. Sci. Rep. 6, 34220–34214. 10.1038/srep34220.

41. Nielsen, M.M., Trolle, C., Vang, S., Hornshoj, H., Skakkebaek, A., Hedegaard, J., Nordentoft, I., Pedersen, J.S., and Gravholt, C.H. (2020). Epigenetic and transcriptomic consequences of excess X-chromosome material in 47,XXX syndrome-A comparison with Turner syndrome and 46,XX females. Am. J. Med. Genet. C: Semin. Med. Genet. 184, 279–293. 10.1002/ajmg.c.31799.

42. Viuff, M., Skakkebaek, A., Johannsen, E.B., Chang, S., Pedersen, S.B., Lauritsen, K.M., Pedersen, M.G.B., Trolle, C., Just, J., and Gravholt, C.H. (2023). X chromosome dosage and the genetic impact across human tissues. Genome Med 15, 21. 10.1186/s13073-023-01169-4.

43. Liu, S., Akula, N., Reardon, P.K., Russ, J., Torres, E., Clasen, L.S., Blumenthal, J., Lalonde, F., McMahon, F.J., Szele, F., et al. (2023). Aneuploidy effects on human gene expression across three cell types. Proc Natl Acad Sci U S A 120, e2218478120. 10.1073/pnas.2218478120.

44. Hughes, J.F., and Page, D.C. (2015). The Biology and Evolution of Mammalian Y Chromosomes. Annu. Rev. Genet. 49, 507–527. 10.1146/annurev-genet-112414-055311.

45. Bellott, D.W., and Page, D.C. (2021). Dosage-sensitive functions in embryonic development drove the survival of genes on sex-specific chromosomes in snakes, birds, and mammals. Genome Res. 10.1101/gr.268516.120.

46. Godfrey, A.K., Naqvi, S., Chmatal, L., Chick, J.M., Mitchell, R.N., Gygi, S.P., Skaletsky, H., and Page, D.C. (2020). Quantitative analysis of Y-Chromosome gene expression across 36 human tissues. Genome Res. 10.1101/gr.261248.120.

47. Pruitt, K.D., Harrow, J., Harte, R.A., Wallin, C., Diekhans, M., Maglott, D.R., Searle, S., Farrell, C.M., Loveland, J.E., Ruef, B.J., et al. (2009). The consensus coding sequence (CCDS) project: Identifying a common protein-coding gene set for the human and mouse genomes. Genome Res. 19, 1316–1323. 10.1101/gr.080531.108.

48. Bray, N.L., Pimentel, H., Melsted, P., and Pachter, L. (2016). Near-optimal probabilistic RNA-seq quantification. Nat Biotechnol 34, 525–527. 10.1038/nbt.3519.

49. Soneson, C., Love, M.I., and Robinson, M.D. (2015). Differential analyses for RNA-seq: transcript-level estimates improve gene-level inferences. F1000Res 4, 1521. 10.12688/f1000research.7563.2.

50. Langmead, B., and Salzberg, S.L. (2012). Fast gapped-read alignment with Bowtie 2. Nat. Methods 9, 357–359. 10.1038/nmeth.1923.

51. Zhang, Y., Liu, T., Meyer, C.A., Eeckhoute, J., Johnson, D.S., Bernstein, B.E., Nusbaum, C., Myers, R.M., Brown, M., Li, W., and Liu, X.S. (2008). Model-based analysis of ChIP-Seq (MACS). Genome Biol. 9, R137. 10.1186/gb-2008-9-9-r137.

52. Subramanian, A., Tamayo, P., Mootha, V.K., Mukherjee, S., Ebert, B.L., Gillette, M.A., Paulovich, A., Pomeroy, S.L., Golub, T.R., Lander, E.S., and Mesirov, J.P. (2005). Gene set enrichment analysis: a knowledge-based approach for interpreting genome-wide expression profiles. Proceedings of the National Academy of Sciences 102, 15545–15550. 10.1073/pnas.0506580102.

53. Liberzon, A., Subramanian, A., Pinchback, R., Thorvaldsdottir, H., Tamayo, P., and Mesirov, J.P. (2011). Molecular signatures database (MSigDB) 3.0. Bioinformatics 27, 1739–1740. 10.1093/bioinformatics/btr260.

54. Teitz, L.S., Pyntikova, T., Skaletsky, H., and Page, D.C. (2018). Selection Has Countered High Mutability to Preserve the Ancestral Copy Number of Y Chromosome Amplicons in Diverse Human Lineages. Am. J. Hum. Genet. 103, 261–275. 10.1016/j.ajhg.2018.07.007.

55. Dohm, J.C., Lottaz, C., Borodina, T., and Himmelbauer, H. (2008). Substantial biases in ultra-short read data sets from high-throughput DNA sequencing. Nucleic Acids Res. 36. ARTN e105 10.1093/nar/gkn425.

56. ’t Hoen, P.A., Friedlander, M.R., Almlof, J., Sammeth, M., Pulyakhina, I., Anvar, S.Y., Laros, J.F., Buermans, H.P., Karlberg, O., Brannvall, M., et al. (2013). Reproducibility of high-throughput mRNA and small RNA sequencing across laboratories. Nat. Biotechnol. 31, 1015–1022. 10.1038/nbt.2702.

57. Lange, J., Skaletsky, H., van Daalen, S.K., Embry, S.L., Korver, C.M., Brown, L.G., Oates, R.D., Silber, S., Repping, S., and Page, D.C. (2009). Isodicentric Y chromosomes and sex disorders as byproducts of homologous recombination that maintains palindromes. Cell 138, 855–869. 10.1016/j.cell.2009.07.042.

58. Lange, J., Noordam, M.J., van Daalen, S.K., Skaletsky, H., Clark, B.A., Macville, M.V., Page, D.C., and Repping, S. (2013). Intrachromosomal homologous recombination between inverted amplicons on opposing Y-chromosome arms. Genomics 102, 257–264. 10.1016/j.ygeno.2013.04.018.

59. Ritchie, M.E., Phipson, B., Wu, D., Hu, Y., Law, C.W., Shi, W., and Smyth, G.K. (2015). limma powers differential expression analyses for RNA-sequencing and microarray studies. Nucleic Acids Res. 43, e47. 10.1093/nar/gkv007.

60. Metsalu, T., and Vilo, J. (2015). ClustVis: a web tool for visualizing clustering of multivariate data using Principal Component Analysis and heatmap. Nucleic Acids Res. 43, W566–570. 10.1093/nar/gkv468.

61. Schiebel, K., Winkelmann, M., Mertz, A., Xu, X., Page, D.C., Weil, D., Petit, C., and Rappold, G.A. (1997). Abnormal XY interchange between a novel isolated protein kinase gene, PRKY, and its homologue, PRKX, accounts for one third of all (Y+)XX males and (Y-)XY females. Hum. Mol. Genet. 6, 1985–1989.

62. Heinz, S., Benner, C., Spann, N., Bertolino, E., Lin, Y.C., Laslo, P., Cheng, J.X., Murre, C., Singh, H., and Glass, C.K. (2010). Simple combinations of lineage-determining transcription factors prime cis-regulatory elements required for macrophage and B cell identities. Mol. Cell 38, 576–589. 10.1016/j.molcel.2010.05.004.

63. Quinlan, A.R., and Hall, I.M. (2010). BEDTools: a flexible suite of utilities for comparing genomic features. Bioinformatics 26, 841–842. 10.1093/bioinformatics/btq033.

64. Chen, B., Gilbert, L.A., Cimini, B.A., Schnitzbauer, J., Zhang, W., Li, G.-W., Park, J., Blackburn, E.H., Weissman, J.S., Qi, L.S., and Huang, B. (2013). Dynamic Imaging of Genomic Loci in Living Human Cells by an Optimized CRISPR/Cas System. Cell 155, 1479–1491. 10.1016/j.cell.2013.12.001.

65. Fulco, C.P., Munschauer, M., Anyoha, R., Munson, G., Grossman, S.R., Perez, E.M., Kane, M., Cleary, B., Lander, E.S., and Engreitz, J.M. (2016). Systematic mapping of functional enhancer–promoter connections with CRISPR interference. Science 354, 769–773. 10.1126/science.aag2445.

66. Stewart, S.A., Dykxhoorn, D.M., Palliser, D., Mizuno, H., Yu, E.Y., An, D.S., Sabatini, D.M., Chen, I.S., Hahn, W.C., Sharp, P.A., et al. (2003). Lentivirus-delivered stable gene silencing by RNAi in primary cells. RNA 9, 493–501. 10.1261/rna.2192803.

67. Horlbeck, M.A., Gilbert, L.A., Villalta, J.E., Adamson, B., Pak, R.A., Chen, Y., Fields, A.P., Park, C.Y., Corn, J.E., Kampmann, M., and Weissman, J.S. (2016). Compact and highly active next-generation libraries for CRISPR-mediated gene repression and activation. eLife 5, e19760. 10.7554/eLife.19760.

68. Price, C., Gill, S., Ho, Z.V., Davidson, S.M., Merkel, E., McFarland, J.M., Leung, L., Tang, A., Kost-Alimova, M., Tsherniak, A., et al. (2019). Genome-Wide Interrogation of Human Cancers Identifies EGLN1 Dependency in Clear Cell Ovarian Cancers. Cancer Res. 79, 2564–2579. 10.1158/0008-5472.CAN-18-2674.

69. R Development Core Team (2020). R: A language and environment for statistical computing (R Foundation for Statistical Computing).

