## Supplemental Information for "The human Y and inactive X chromosomes similarly modulate autosomal gene expression"

**Figures S1-S19**

**Tables S1-S12** *(provided as separate .xlsx files)*

**Figure S1**

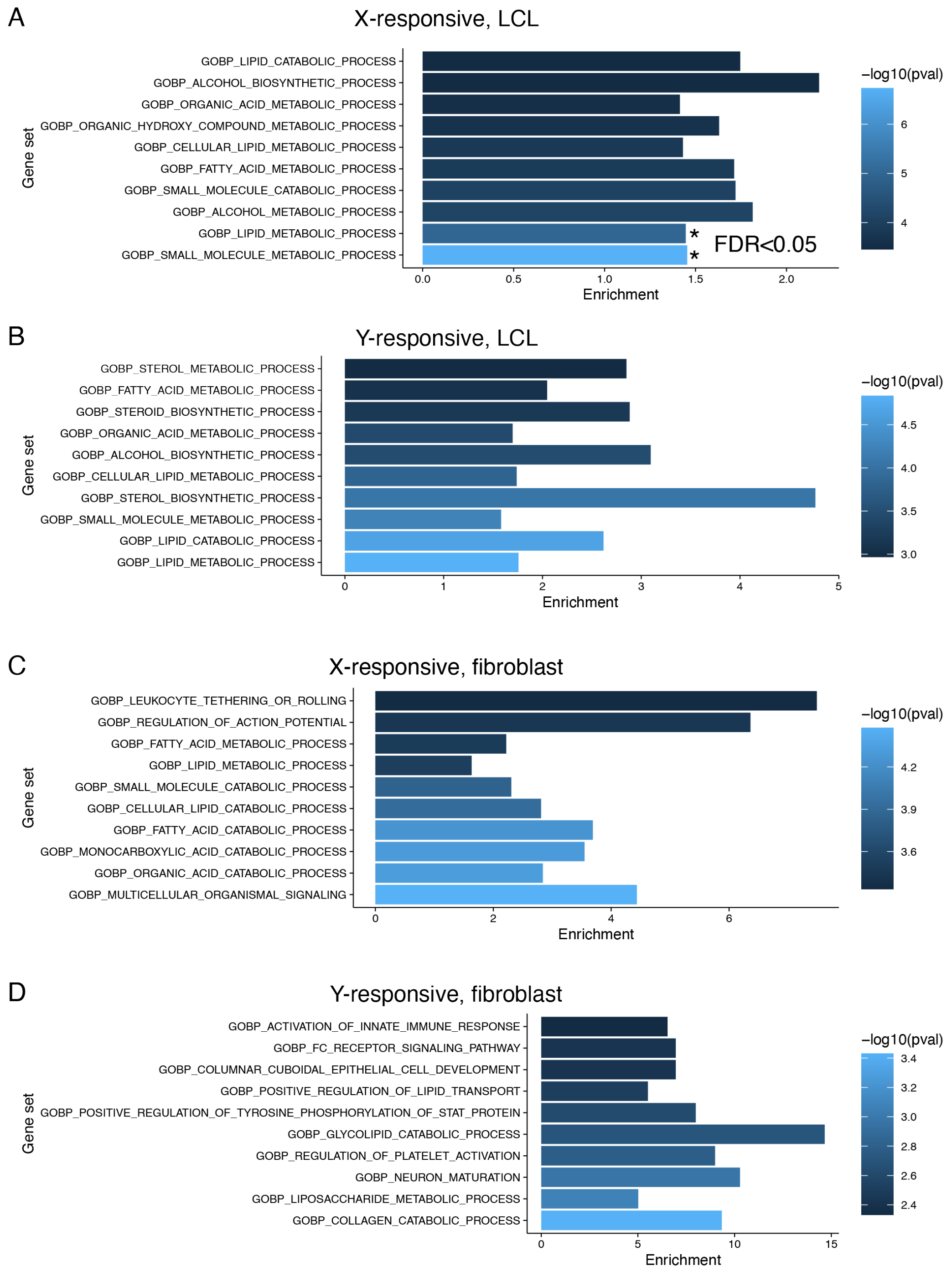


**Figure S1. Analysis of gene ontology category enrichment in autosomal genes that significantly respond to Chr X or Y copy number, related to Figure 1.** Enrichment of

“Biological Process” Gene Ontology categories was performed separately for autosomal genes that significantly responded to Chr X or Y copy number in LCLs or fibroblasts. All expressed genes in LCLs or fibroblasts were used as the background set. Shown here are the top 10 categories based on P value. Stars indicate those that were below the Benjamini-Hochberg-corrected P value (FDR) of 0.05. There were no significant categories (by FDR threshold) for genes that responded to Chr Y copy number in LCLs, nor for genes that responded to Chr X or Y copy number in fibroblasts.

**Figure S2**

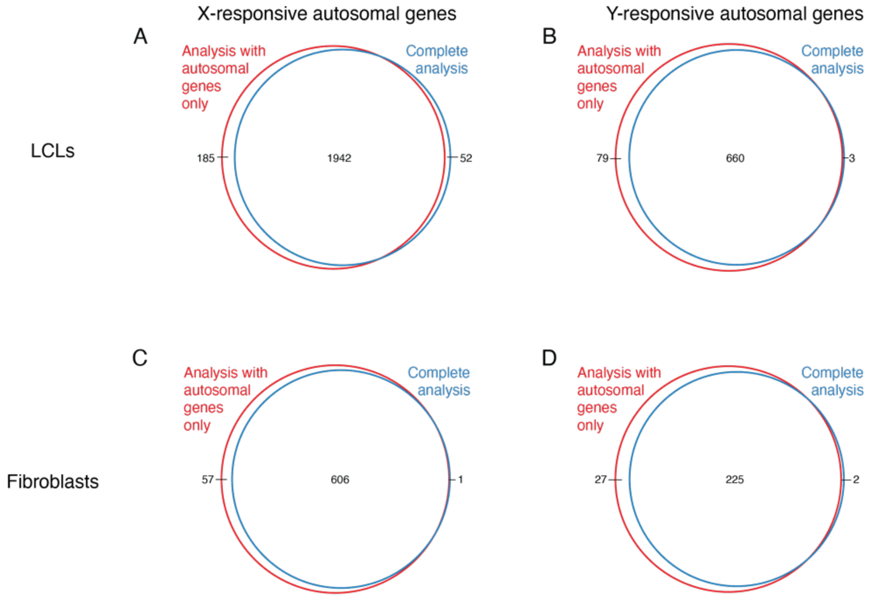


**Figure S2. Mapping to a transcriptome with or without the sex chromosomes does not affect X- and Y-responsive autosomal genes in LCLs and fibroblasts, related to Figure 1.** Read depth normalization between samples and differential expression was performed using an annotation only containing autosomal genes to rule out artifacts of bulk over-expression of sex chromosome genes, and compared to the analysis conducted using the complete annotation. Significantly X-responsive genes in LCLs (A) and fibroblasts (B) were almost completely overlapping between the two analyses, as were the significantly Y-responsive genes in LCLs (C) and fibroblasts (D).

**Figure S3**

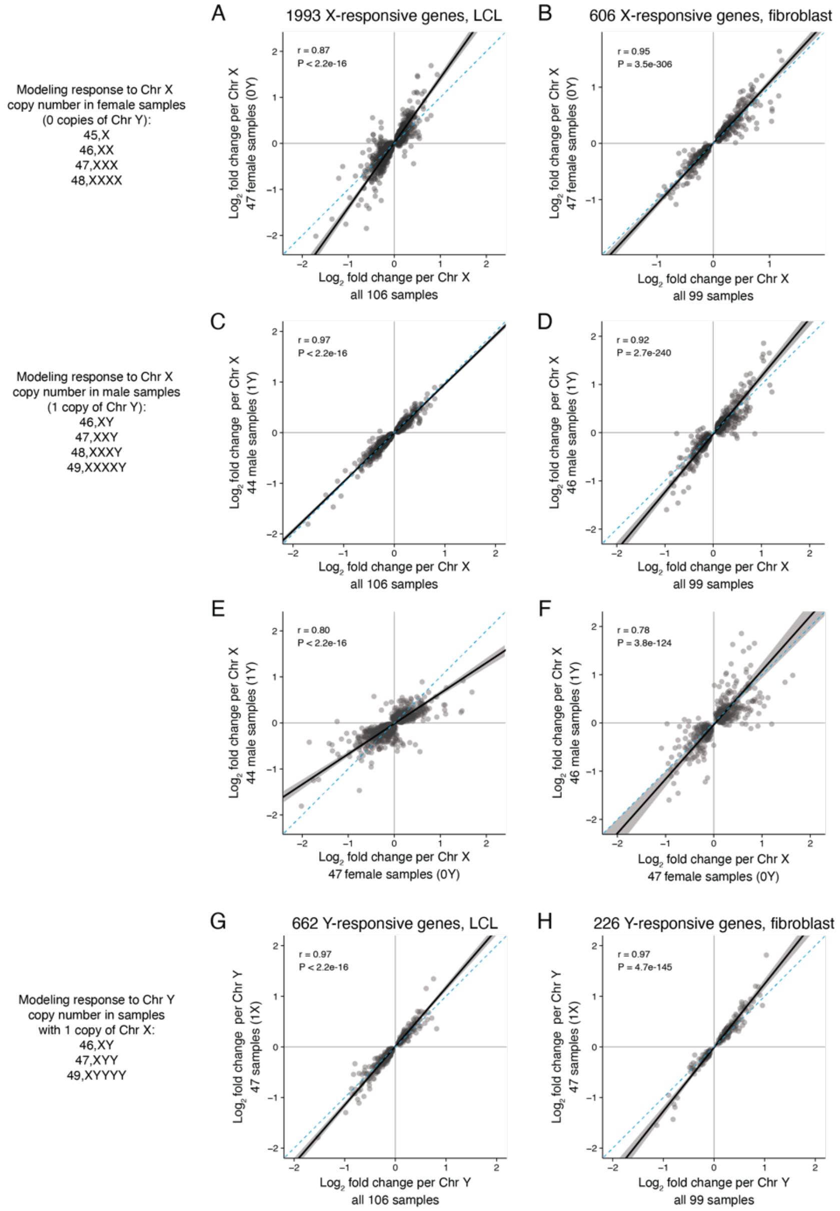


**Figure S3. Modeling Chr X or Y copy number separately is highly correlated with models including both covariates, related to Figure 1. (A-D)** Scatterplots comparing models of gene expression as a function of Chr X copy number in female samples (zero Y chromosomes; [A, B]) or male samples with one Y chromosome (C, D) with the full model show high correlations. (**E, F)** Scatterplots comparing models of gene expression as a function of Chr X copy number in male samples with one Y chromosome and female samples (no Y chromosomes) show high correlations. **(G, H)** Scatterplots comparing a model of gene expression as a function of Chr Y copy number in male samples with one X chromosome with the full model reveal a high correlation. Black line and grey shading, Deming regression and 95% confidence interval; blue dashed line, identity (X=Y) line. Pearson correlation coefficients and P values are indicated.

**Figure S4**

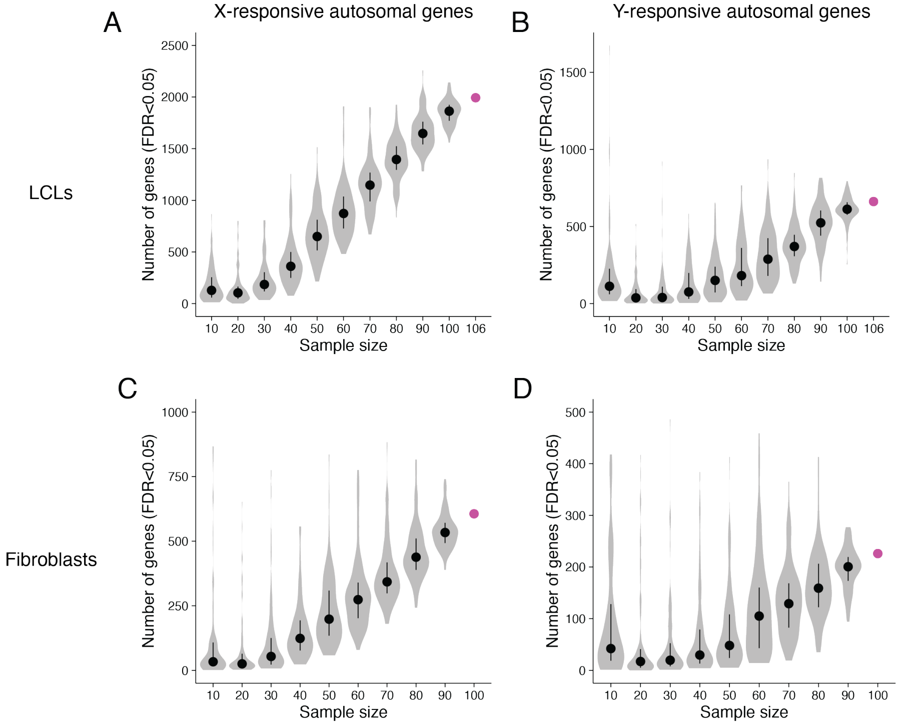


**Figure S4. Detection of autosomal genes significantly responsive to Chr X and Y copy number is not saturated at the current sample size, related to Figure 1.** To determine whether we had saturated the signal of significantly Chr X- or Y-responsive genes at the given sample size, we performed a bootstrapping analysis. Violin plots with median (dots) and interquartile range (whiskers) showing the number of Chr X-responsive (**A, C**) or Y-responsive (**B, D**) autosomal genes detected in random subsets of the LCL (A,B) or fibroblast (C,D) RNA-seq libraries. The number of samples in each subset is shown on the x-axis. 100 random samples were obtained at each sample size. Pink dots show the number of Chr X- and Y-responsive genes detected in the full set of samples. The large number of genes detected in some of the smallest subsets is caused by unbalanced karyotype distributions.

#### Figure S5

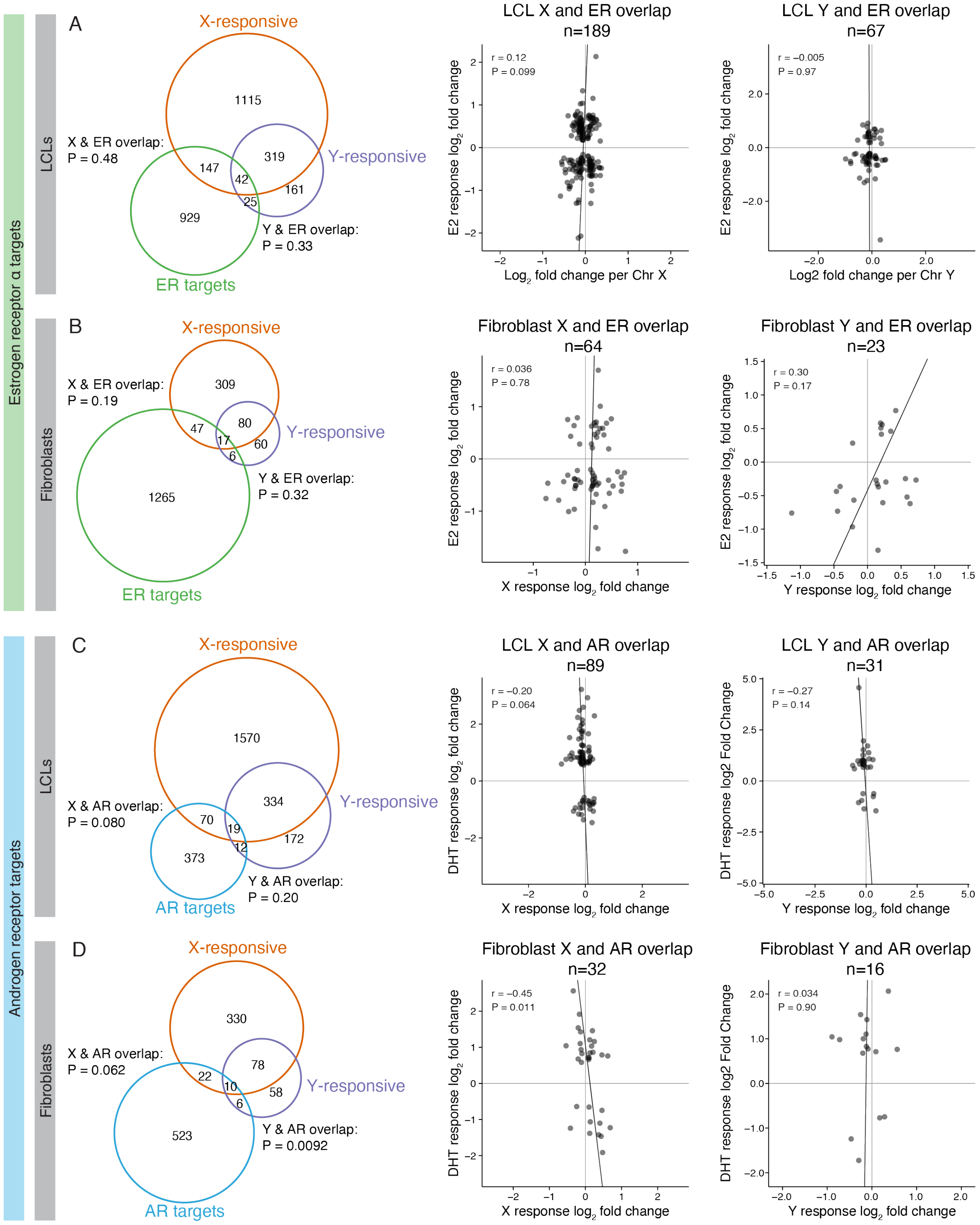


**Figure S5. The response to Chr X- and Y-responsive genes is distinct from the response to sex hormones, as defined by estrogen receptor alpha and androgen receptor direct target**

**genes, related to Figure 1. (A-B)** *Left,* Venn diagrams showing overlap of ERα direct target genes (bound by ERα within 30 kb of the transcription start site and differentially expressed [$P_{adj}$<0.05] with 17-ß estradiol [E2] treatment) and Chr X- or Y-responsive genes in LCLs (A) or fibroblasts (B). Venn diagrams were restricted to autosomal genes expressed in both MCF7 cells (where ERα direct targets were defined) and LCLs or fibroblasts. *Right,* scatterplots of overlapping genes showing the relationship between response to E2 and Chr X or Y copy number. **(C-D)** *Left,* Venn diagrams showing overlap of AR direct target genes (bound by AR within 30 kb of the transcription start site and differentially expressed [$P_{adj}$<0.05] with dihydrotestosterone [DHT] treatment) and Chr X- or Y-responsive genes in LCLs (C) or fibroblasts (D). Venn diagrams were restricted to autosomal genes expressed in both LNCaP cells (where AR direct targets were defined) and LCLs or fibroblasts. *Right,* scatterplots of overlapping genes showing the relationship between response to DHT and Chr X or Y copy number. Venn diagram P values, hypergeometric test; scatterplots show Deming regressions (black lines), Pearson correlation coefficients and P values.

**Figure S6**

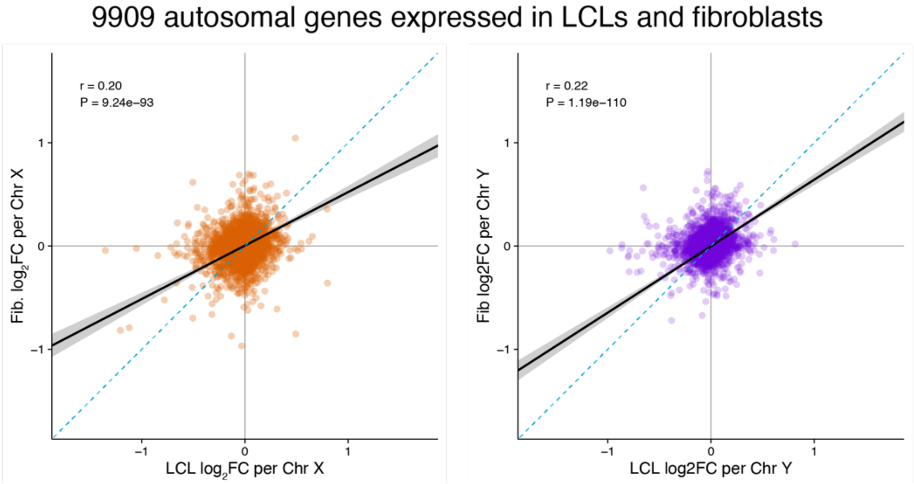


**Figure S6. Response to Chr X or Y copy number is weakly correlated between cell types, related to Figure 1.** Scatterplots comparing the effect sizes of the response to Chr X (left) or Y (right) copy number in LCLs versus fibroblasts for all autosomal genes expressed in both cell types. Black line and grey shading, weighted Deming regression and 95% confidence interval; blue dashed line, identity (X=Y) line; Pearson correlation coefficients and P values are indicated.

**Figure S7**

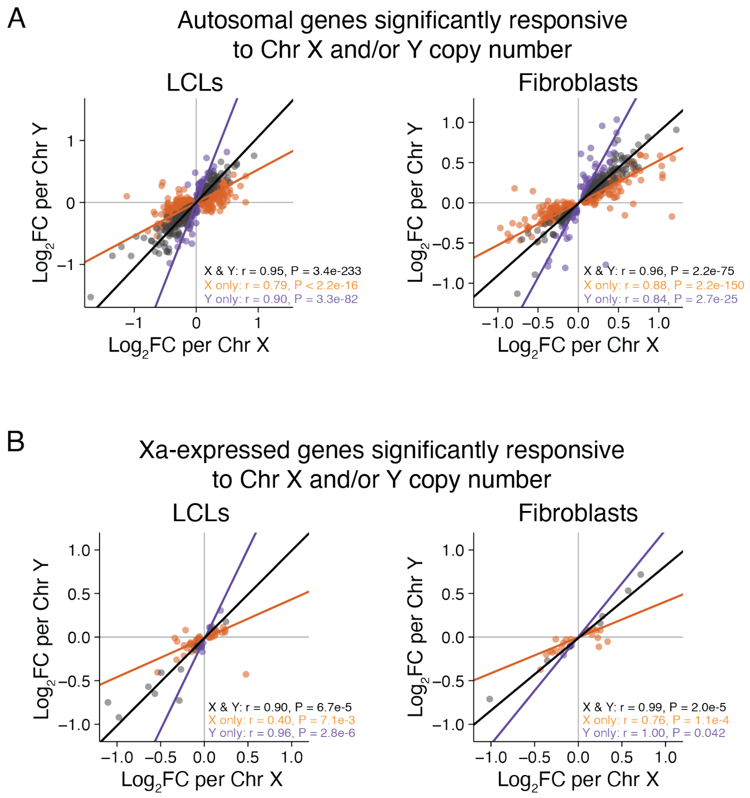


**Figure S7. Correlation plots for significantly X and/or Y responsive genes, related to Figure 2.** Scatterplots showing the relationship between the effects per additional copy of Chr X or Y for autosomal genes **(A)** or Xa-expressed genes **(B)** called as significantly responsive ($P_{adj}$<0.05) to either Chr X, Chr Y, or both. Weighted Deming regression lines, Pearson correlation coefficients, and P values of the correlation are included in each plot.

**Figure S8**

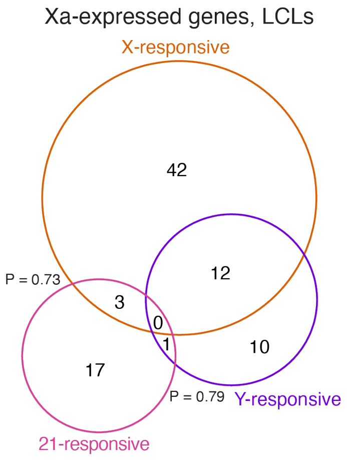


**Figure S8. For Xa-expressed genes, the response to Chr 21 copy number is distinct from the shared response to Chr X and Chr Y copy number, related to Figure 3.** Venn diagram of Xa-expressed genes that significantly respond to Chr 21, Chr X, or Chr Y copy number. P values, hypergeometric test.

**Figure S9**

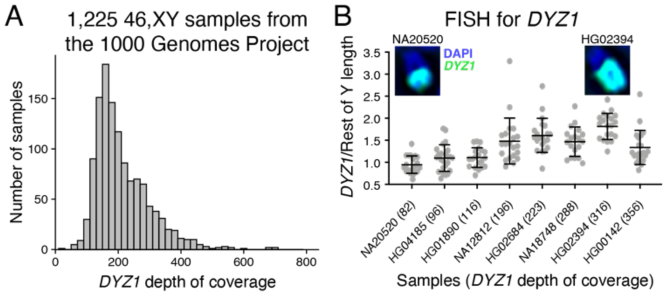


**Figure S9. Wide range of Y heterochromatin lengths in males from the 1000 Genomes Project, related to Figure 4.** (A) Histogram of *DYZ1* depth of coverage – a proxy for Chr Y heterochromatin length – for 1,225 46,XY samples from the 1000 genomes project. (B) Quantification of DNA fluorescent in situ hybridization (FISH) signal for *DYZ1* on Y chromosome spreads from 1000 Genomes Project samples. Each point represents the ratio of the length of the Chr Y segment marked by *DYZ1* probes, compared to the length of the Chr Y segment not marked by *DYZ1* probes in mitotic chromosome spreads from 8 individuals in the 1000 genomes project. Inset, representative images from the sample with the smallest and largest *DYZ1*-marked to non-*DYZ1*-marked Y chromosome ratios.

### Figure S10

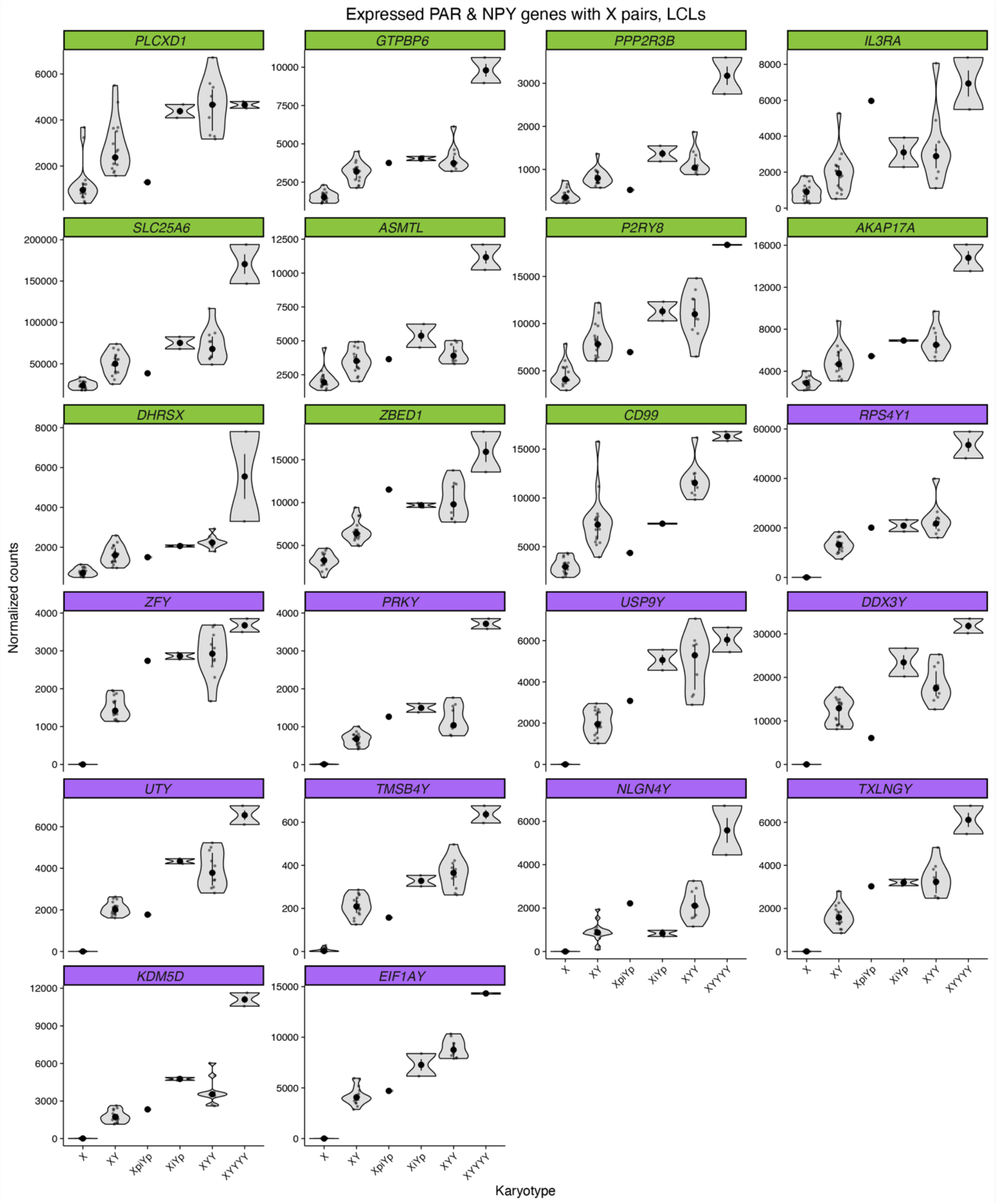
Expressed PAR & NPY genes with X pairs, LCLs
PLCXD1
GTPBP6
PPP2R3B
IL3RA
SLC25A6
ASMTL
P2RY8
AKAP17A
DHRSX
ZBED1
CD99
RPS4Y1
ZFY
PRKY
USP9Y
DDX3Y
UTY
TMSB4Y
NLGN4Y
TXLNGY
KDM5D
EIF1AY
Normalized counts
X
XY
XpiYp
XiYp
XYY
XYYYY
Karyotype

**Figure S10. Chr Y gene expression in samples with variant Y chromosomes, related to Figure 4.** Violin plots showing median and interquartile range for all expressed Chr Y genes (PAR1 and NPY) for samples with two types of variant Y chromosomes: isoYp (XiYp) with recombination in the P1 palindrome, pseudoisoYp (XpiYp) with recombination between IR4 inverted repeats, or samples with zero to four copies of Chr Y and one copy of Chr X. Green, PAR1 genes; purple, NPY genes with pairs on Chr X.

**Figure S11**

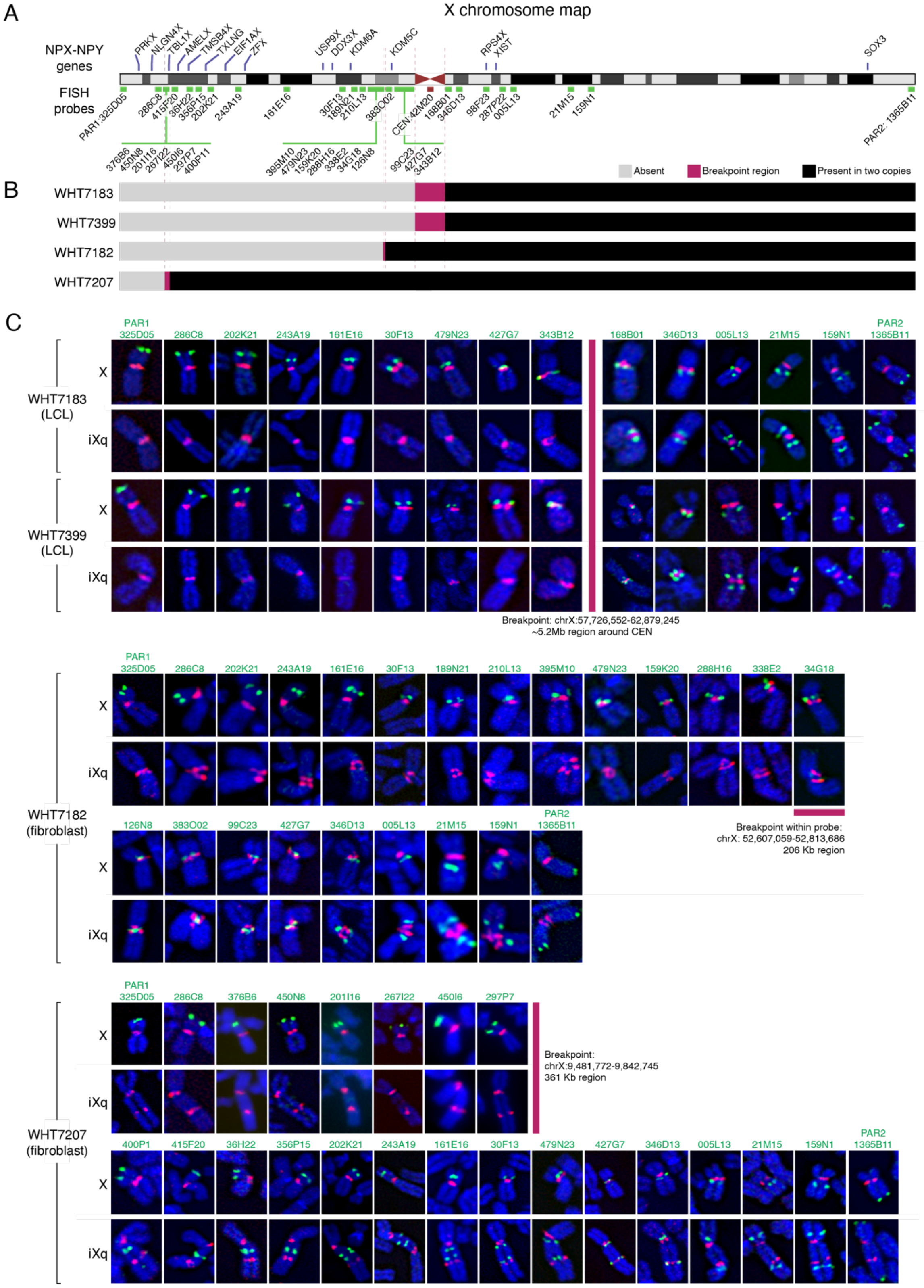
A
X chromosome map
NPX-NPY genes
FISH probes
B
Absent
Breakpoint region
Present in two copies
WHT7183
WHT7399
WHT7182
WHT7207
C
WHT7183 (LCL)
WHT7399 (LCL)
X
iXq
Breakpoint: chrX:57,726,552-62,879,245
~5.2Mb region around CEN
WHT7182 (fibroblast)
Breakpoint within probe:
chrX: 52,607,059-52,813,686
206 Kb region
WHT7207 (fibroblast)
Breakpoint:
chrX:9,481,772-9,842,745
361 Kb region

**Figure S11. Determination of X isochromosome breakpoints in four individuals using DNA FISH, related to Figure 5. (A)** Chr X ideogram showing locations of NPX-NPY gene pairs and FISH probes. The centromere probe 42M20 was used in all experiments as a reference (in red), while all other probes were labelled in green. **(B)** Locations of breakpoints on X isochromosomes as determined by FISH. The locations of the breakpoints correspond to the schematic of the X Chr in (A). Grey, region is absent in iXq; pink, breakpoint region; black, region is present in two copies in iXq. **(C)** Breakpoints for iXq chromosomes were refined using DNA FISH. For each individual and probe, an image of a normal Chr X and iXq are included from the same chromosome spread for comparison. Images for WHT7183 and WHT7399 were obtained from FISH on spreads from LCLs, while WHT7182 and WHT7202 were obtained from FISH on spreads from fibroblasts.

#### Figure S12

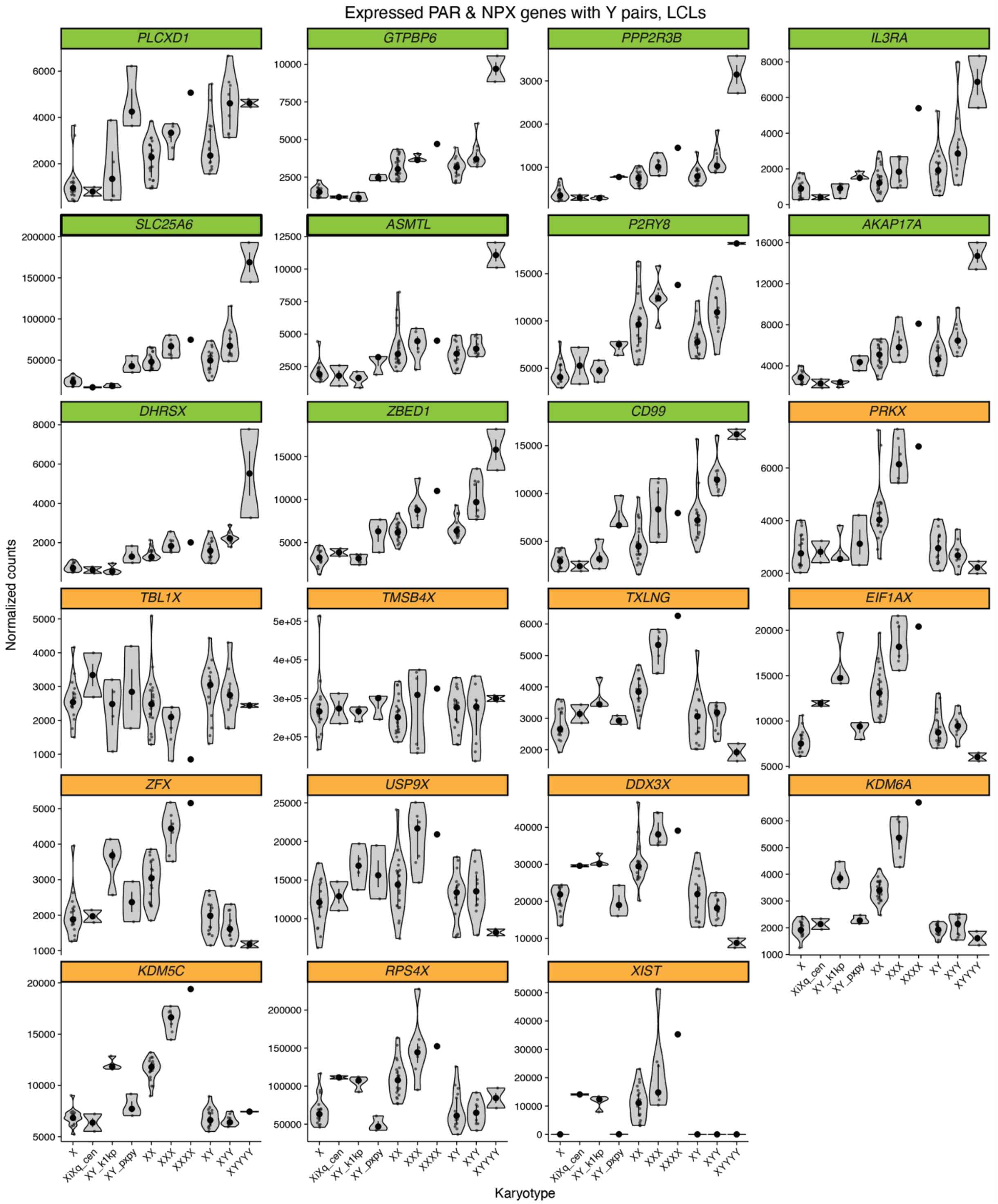
Expressed PAR & NPX genes with Y pairs, LCLs
PLCXD1
GTPBP6
PPP2R3B
IL3RA
SLC25A6
ASMTL
P2RY8
AKAP17A
DHRSX
ZBED1
CD99
PRKX
TBL1X
TMSB4X
TXLNG
EIF1AX
ZFX
USP9X
DDX3X
KDM6A
KDM5C
RPS4X
XIST
Normalized counts
Karyotype
X
XiXq_cen
XY_k1kp
XY_pxpy
XX
XXX
XXXX
XY
XYY
XYYYY

**Figure S12. Expression levels of PAR1 and NPX genes with NPY homologs in LCL samples with structural variants of Chr X or Chr Y, related to Figure 5.** Violin plots showing median and interquartile range for expressed PAR1 genes (green) and NPX genes with NPY homologs and *XIST* (orange) for samples with Chr X or Y structural variants or samples with one to four copies of Chr X and one to four copies of Chr Y. XiXq_cen: 46,X,i(Xq) samples with recombination in the centromere; XY_k1kp: 46,X,t(X;Y)(*ANOS1-ANOS2P*); XY_pxpy: 46,X,t(X;Y)(*PRKX-PRKY*).

**Figure S13**

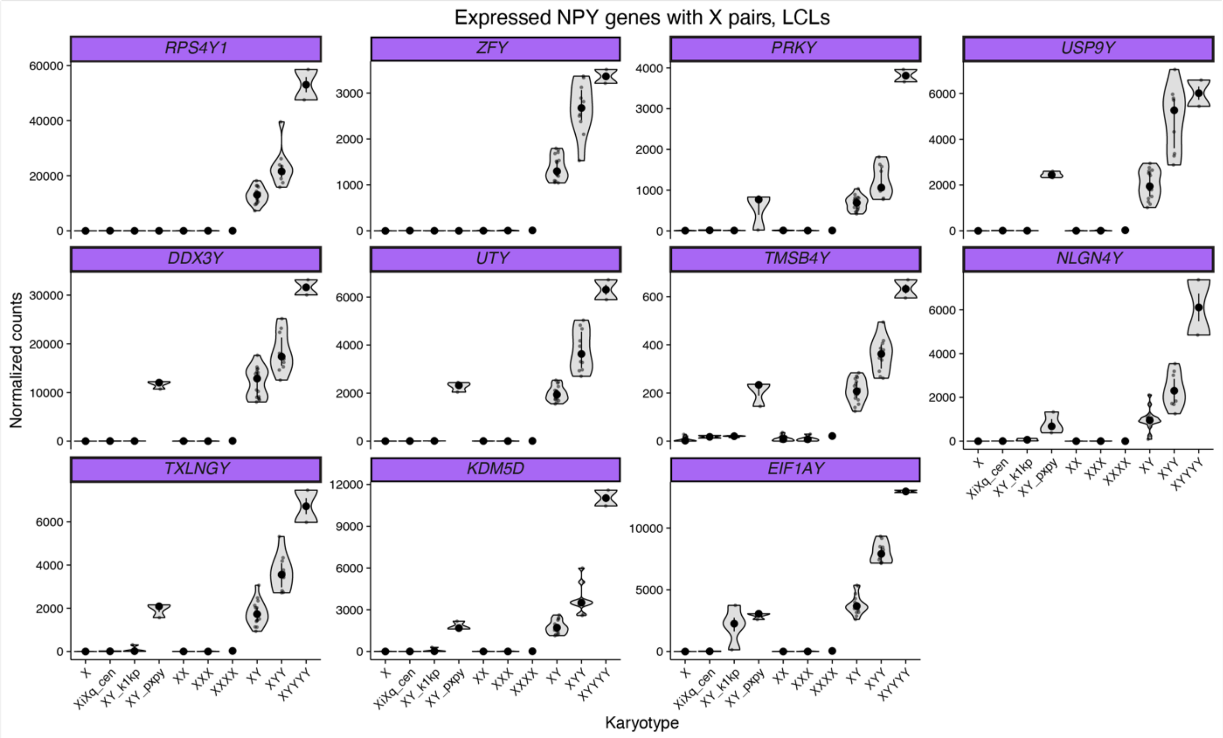


**Figure S13. Expression levels of NPY genes with NPX homologs in LCL samples with structural variants of Chr X or Chr Y, related to Figure 5.** Violin plots showing median and interquartile range for expressed NPY genes for samples with Chr X or Y structural variants or samples with one to four copies of Chr X and one to four copies of Chr Y. XiXq_cen: 46,X,i(Xq) samples with recombination in the centromere; XY_k1kp: 46,X,t(X;Y)(*ANOS1-ANOS2P*); XY_pxpy: 46,X,t(X;Y)(*PRKX-PRKY*).

#### Figure S14

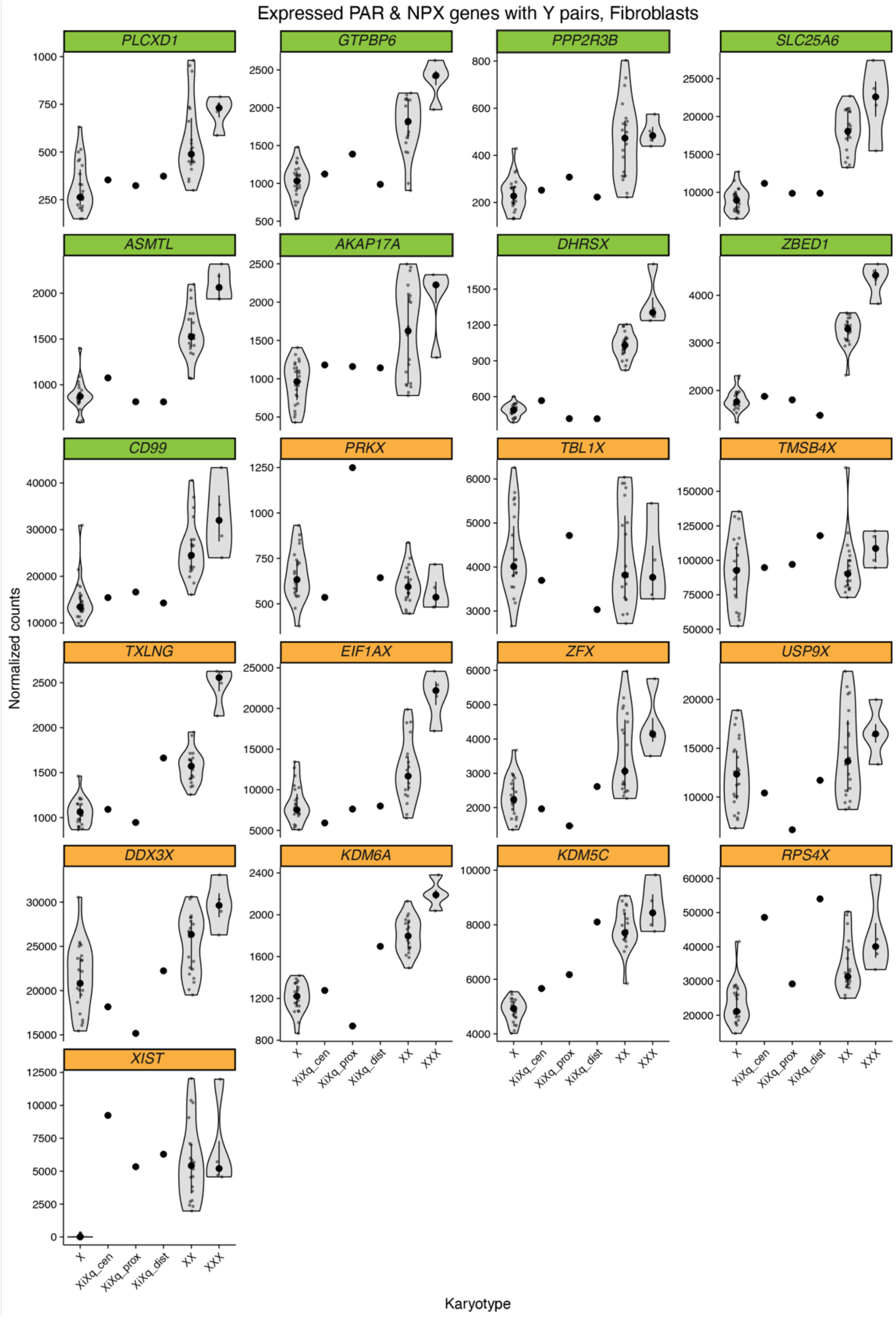
Expressed PAR & NPX genes with Y pairs, Fibroblasts
PLCXD1
GTPBP6
PPP2R3B
SLC25A6
ASMTL
AKAP17A
DHRSX
ZBED1
CD99
PRKX
TBL1X
TMSB4X
TXLNG
EIF1AX
ZFX
USP9X
DDX3X
KDM6A
KDM5C
RPS4X
XIST
Normalized counts
X
XiXq_cen
XiXq_prox
XiXq_dist
XX
XXX
Karyotype

**Figure S14. Expression levels of PAR1 and NPX genes with NPY homologs in fibroblast samples with structural variants of Chr X, related to Figure 5.** Violin plots showing median and interquartile range for expressed PAR1 genes (green) and NPX genes with NPY homologs and *XIST* (orange) for samples with Chr X structural variants or samples with one to three copies of Chr X. XiXq_cen: 46,X,i(Xq) samples with recombination in the centromere; XiXq_prox: 46,X,i(Xq) samples with recombination in proximal Xp; XiXq_dist: 46,Xi(Xq) samples with recombination in distal Xp.

**Figure S15**

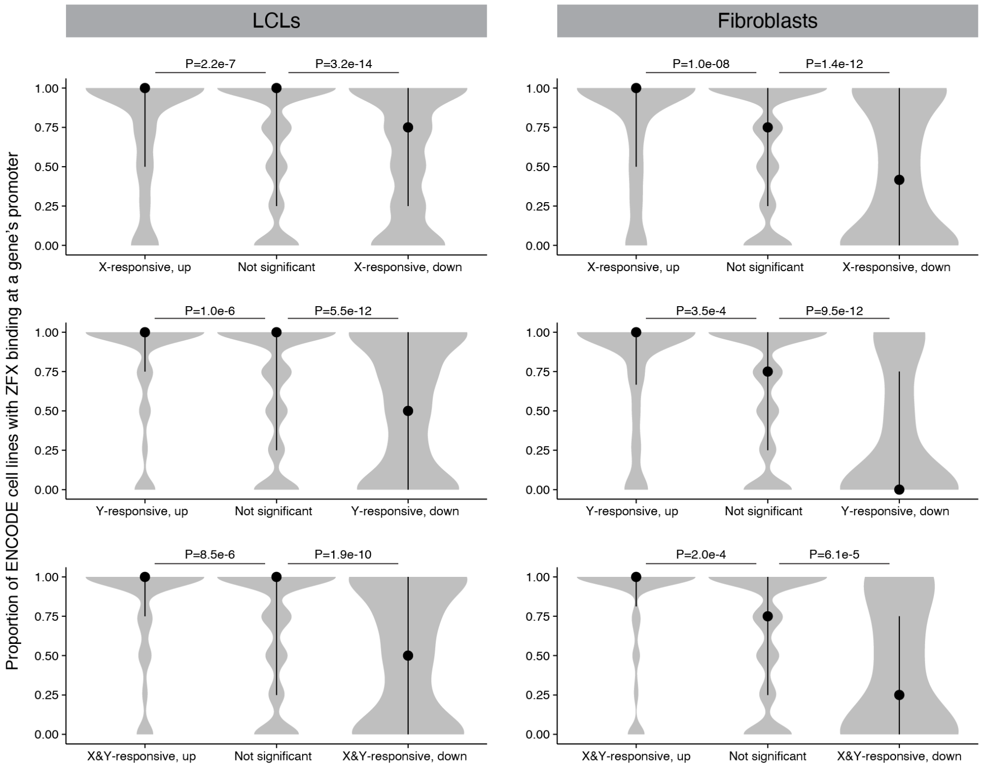


**Figure S15. ZFX protein binding is enriched in promoters of autosomal genes whose expression increases in response to Chr X and/or Y copy number, and is depleted in promoters of genes whose expression decreases.** For autosomal genes with or without significant expression changes with Chr X or Chr Y copy number, we investigated ZFX binding using publicly available ChIP-seq data from four cell lines. Plotted is the proportion of cell lines with ZFX binding at the gene's promoter (in cell lines where the gene is expressed). For example, a gene expressed in three of the four cell types with a ZFX peak in the promoter in those three cell types would have a proportion bound of 1. P values, Wilcoxon rank sum test.

#### Figure S16

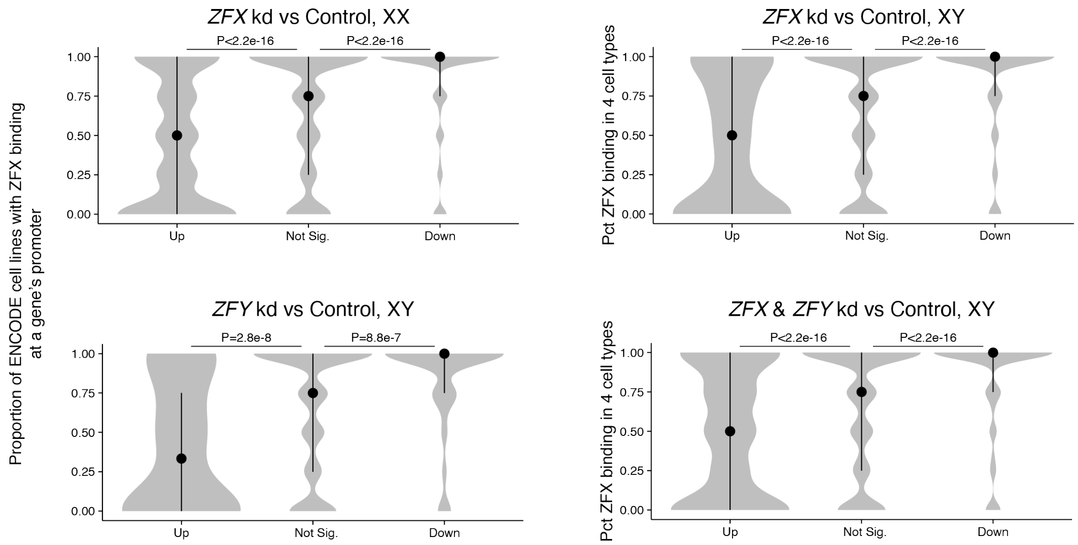


**Figure S16. ZFX protein binding is enriched in promoters of autosomal genes that decrease in response to *ZFX* and/or *ZFY* knockdown in fibroblasts, and is depleted in promoters of genes that increase.** For autosomal genes with or without significant expression changes in each of the four CRISPRi experiments, we investigated ZFX binding using publicly available ChIP-seq data from four cell lines (as in **Fig. S15**). Plotted is the proportion of cell lines with ZFX binding at the gene's promoter (in cell lines where the gene is expressed). P values, Wilcoxon rank sum test.

**Figure S17**

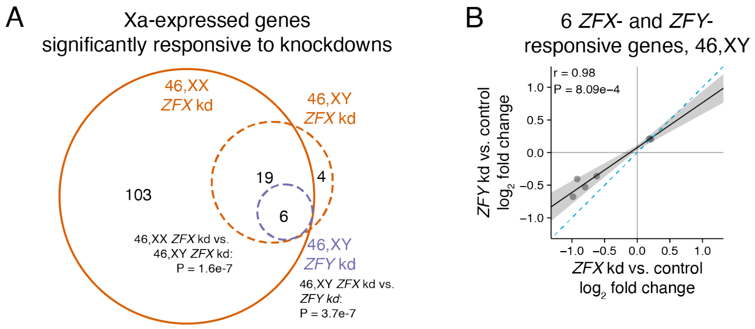


**Figure S17. All *ZFY*-responsive Xa-expressed genes are also *ZFX*-responsive.** (A) Venn diagram of Xa-expressed genes significantly differentially expressed upon knockdown of *ZFX* in XX or XY cells or upon knockdown of *ZFY* in XY cells. P values, hypergeometric test. (B) Scatterplot of the four significantly *ZFX*- and *ZFY*-responsive Xa-expressed genes in 46,XY cells comparing effects of *ZFX* versus *ZFY* knockdowns. Black line and grey shading, weighted Deming regression and 95% confidence interval; blue dashed line, identity (X=Y) line; Pearson correlation coefficient and P value are indicated.

#### Figure S18

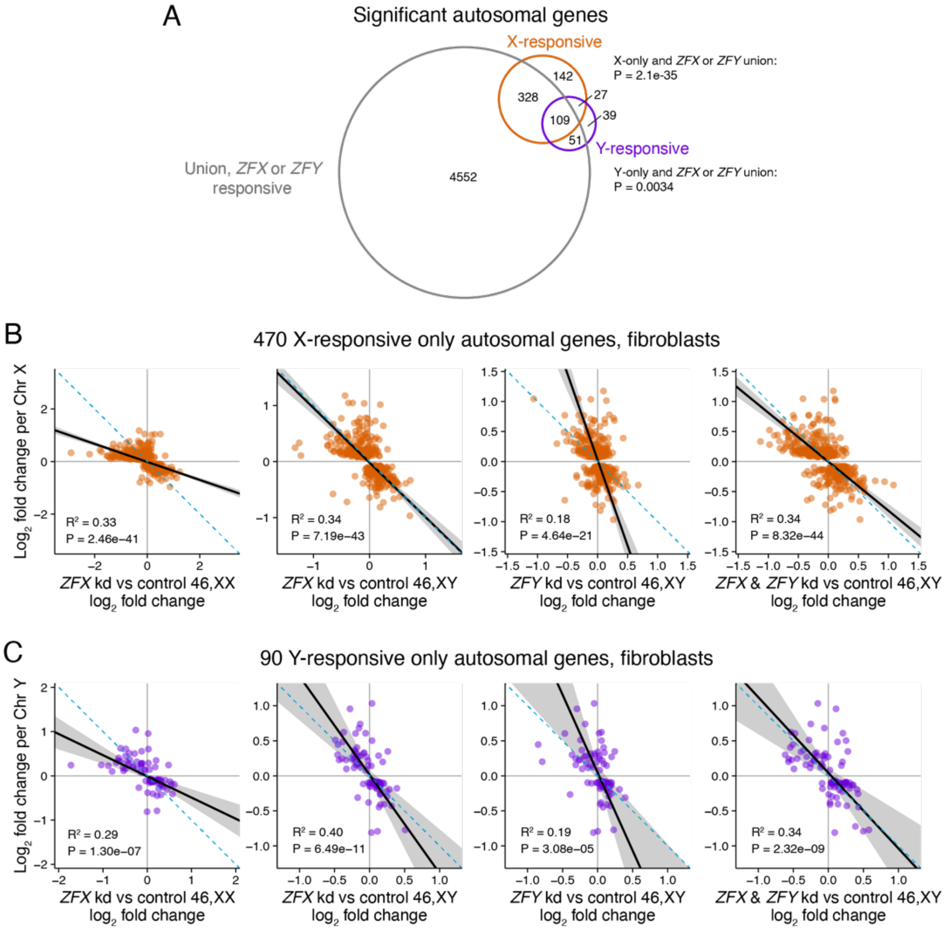


**Figure S18. *ZFX* and *ZFY* response also explains variation in X-responsive-only and Y-responsive-only autosomal genes, related to Figure 7. (A)** Venn diagram of genes significantly responsive to either Chr X or Y copy number in fibroblasts and the union of genes responsive to *ZFX* or *ZFY* across all four CRISPRi experiments. P values, hypergeometric test. **(B-C)** Scatterplots for X-responsive-only (B) or Y-responsive-only (C) genes comparing the effect size in response to Chr X (B) or Chr Y (C) copy number with the effect size in response to four *ZFX* or *ZFY* CRISPRi experiments. Black line and grey shading, weighted Deming regression and 95% confidence interval; blue dashed line, identity line (X=Y). Coefficients of determination and P values are indicated.

**Figure S19**

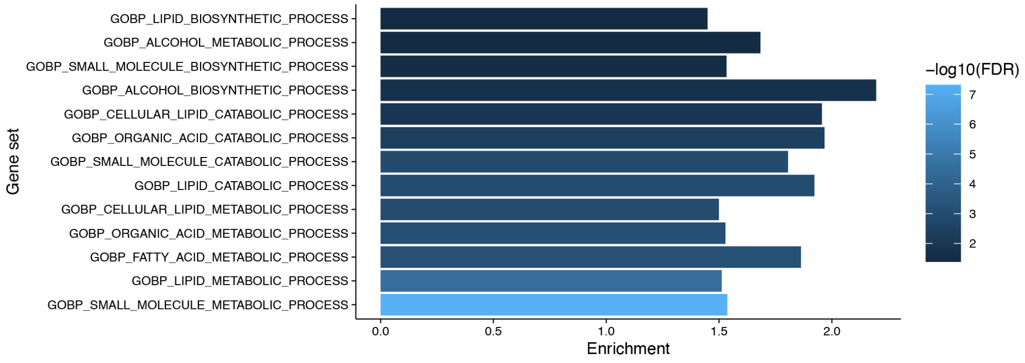


**Figure S19. Functional enrichment in union of significant Chr X or Y responsive autosomal and Xa-expressed genes in LCLs and fibroblasts.** Enrichment of "Biological Process" Gene Ontology categories was performed for the union of autosomal and Xa-expressed genes genes that significantly responded to Chr X or Y copy number in LCLs and fibroblasts. The union of all expressed autosomal and Xa-expressed genes in LCLs and fibroblasts was used as the background set. Shown here are all categories that fell below the FDR threshold of 0.05.

#### SUPPLEMENTAL TABLES

**Table S1.** Metadata for RNA-seq of LCLs or fibroblasts from individuals with naturally-occurring variations in chromosome copy number or structure, related to Figures 1, 3, 4, and 5

**Table S2.** Expression change per copy of Chr X or Chr Y for expressed autosomal genes in LCLs and fibroblasts, related to Figure 1

**Table S3.** Reanalysis of publicly available data to identify AR and ER direct target genes, related to Figure 1

**Table S4.** Genes expressed from the active X chromosome only (silenced on Xi) that significantly change in expression in response to Chr X and Chr Y copy number, related to Figure 2

**Table S5.** Expression change per copy of Chr 21 for expressed autosomal genes in LCLs, related to Figure 3

**Table S6.** Differential gene expression analysis as a function of *DYZ1* depth of coverage, related to Figure 4

**Table S7.** Sex chromosome structural variant breakpoints and gene content, related to Figures 4 and 5

**Table S8.** Sequence motifs in promoters of Chr X- and Y-responsive genes in LCLs and fibroblasts, related to Figure 6

**Table S9.** Genome-wide binding analysis of ZFX across ENCODE cell lines, related to Figure 6

**Table S10.** Expression analysis of *ZFX* and *ZFY* CRISPRi knockdowns, related to Figure 6

**Table S11.** Sequence motifs in promoters of genes affected by *ZFX* and *ZFY* knockdowns, related to Figure 6

**Table S12.** Quantification of genome-wide response to Chr X and Y copy number in LCLs and fibroblasts, related to Figures 1 and 2
